# The evolution of egg trading in simultaneous hermaphrodites

**DOI:** 10.1101/460386

**Authors:** Jorge Peña, Georg Nöldeke, Oscar Puebla

**Affiliations:** Institute for Advanced Study in Toulouse, University of Toulouse Capitole, Toulouse, France; Faculty of Business and Economics, University of Basel, Basel, Switzerland; GEOMAR Helmholtz Centre for Ocean Research Kiel, Kiel, Germany; University of Kiel, Faculty of Mathematics and Natural Sciences, Kiel, Germany; Smithsonian Tropical Research Institute, Panamá, República de Panamá

**Keywords:** Egg trading, simultaneous hermaphroditism, cooperation, direct reciprocity

## Abstract

Egg trading, whereby simultaneous hermaphrodites exchange each other’s eggs for fertilization, constitutes one of the few rigorously documented and most widely cited examples of direct reciprocity among unrelated individuals. Yet how egg trading may initially invade a population of non-trading simultaneous hermaphrodites is still unresolved. Here, we address this question with an analytical model that considers mate encounter rates and costs of egg production in a population that may include traders (who provide eggs for fertilization only if their partners also have eggs to reciprocate), providers (who provide eggs regardless of whether their partners have eggs to reciprocate), and withholders (“cheaters” who only mate in the male role and just use their eggs to elicit egg release from traders). Our results indicate that a combination of intermediate mate encounter rates, sufficiently high costs of egg production, and a sufficiently high probability that traders detect withholders (in which case eggs are not provided) is conducive to the evolution of egg trading. Under these conditions traders can invade—and resist invasion from—providers and withholders alike. The prediction that egg trading evolves only under these specific conditions is consistent with the rare occurrence of this mating system among simultaneous hermaphrodites.

## Introduction

Sexual conflict arises when there is a conflict of interest between the two members of a mating pair over sexual reproduction (Hammerstein & Parker, 1987; Kokko & Jennions, 2014). In simultaneous hermaphrodites such a conflict arises with respect to the male and female functions, and often manifests as a preference for mating in the male role (Charnov, 1979). Such preference has been interpreted as a direct consequence of anisogamy: since eggs are more energetically costly to produce than sperm, reproductive success is expected to be limited by access to eggs specifically (Bateman (1948), see also Parker & Birkhead (2013) for a more recent perspective). Mating in the male role should therefore be preferred, which creates a conflict of interest between mating partners: both would prefer to mate in the male role, but for the mating to be successful one partner needs to mate in the less preferred female role (Leonard, 1993).

Egg trading is a specific mating system whereby simultaneous hermaphrodites trade each other’s eggs for fertilization, which contributes to resolve this type of conflict. Egg trading evolved independently in fishes (Fischer, 1980, 1984; Oliver, 1997; Petersen, 1995; Pressley, 1981) and polychaetes (Picchi et al., 2018; Sella, 1985; Sella & Lorenzi, 2000; Sella et al., 1997; Sella & Ramella, 1999). When mating, a pair of egg traders take turns in fertilizing each other’s eggs. By linking male reproductive success to female reproductive success, egg trading disincentivizes spawning in the male role predominantly or exclusively, as opportunities to fertilize a partner’s eggs depend on providing eggs to that partner (Fischer, 1980). More broadly, egg trading constitutes one of the few rigorously documented and most widely cited examples of direct reciprocity among unrelated individuals in animals (Axelrod & Hamilton, 1981). Direct reciprocity (also known as “reciprocal altruism”; Trivers 1971) operates when an individual acts at an immediate fitness cost to benefit another individual, who in turn reciprocates that benefit back. It provides a mechanism for the evolution of cooperation among genetically unrelated individuals (Lehmann & Keller, 2006; Nowak, 2006; Sachs et al., 2004; Van Cleve & Akçay, 2014).

To date, most theoretical work on egg trading has sought to explain (i) its evolutionary stability against invasion by “cheaters” (referred here as “withholders”) who fertilize their partners’ eggs but do not reciprocate by releasing eggs (Crowley & Hart, 2007; Friedman & Hammerstein, 1991; Leonard, 1990), and (ii) its role in making simultaneous hermaphroditism evolutionarily stable relative to gonochorism (Fischer, 1980; Henshaw et al., 2015). While these studies addressed the stability and evolutionary consequences of egg trading once it is already established, how egg trading may evolve in the first place turned out to be a problematic question. Axelrod & Hamilton (1981) speculated that egg trading might have evolved through a low-density phase that would have favored self-fertilization and inbreeding, which would have in turn allowed kin selection to operate. However, this hypothesis has been challenged on the grounds that many egg traders do not (and might not have the physiological ability to) self-fertilize (Fischer, 1981, 1988).

More recently, Henshaw et al. (2014) provided a combination of analytical and simulation models that constitutes the first thorough attempt to explicitly address the evolution of egg trading. Their analytical model considers mate encounters in a population that includes non-traders (individuals who provide eggs at every mating opportunity, referred here as “providers”) and traders (individuals who provide eggs only if their partner have eggs to reciprocate). Their results show that, as with other instances of direct reciprocity (André, 2014), egg trading is under positive frequency-dependent selection and counterselected unless the proportion of traders in the population reaches a critical threshold. Egg trading can therefore only reach fixation in this model when the strategy is already represented by a certain proportion of the population, leaving it open how rare egg-trading mutants may initially persist and spread. Henshaw et al. (2014) showed that the egg-trading invasion barrier is easier to overcome when encounters between mates are frequent, as such high encounter rates increase the chances that a rare egg trader will find a partner with eggs to reciprocate. This relationship between encounter rates and the evolution of egg trading raises an interesting dilemma since high encounter rates have also been found to destabilize egg trading by allowing withholders to invade a population of egg traders (Crowley & Hart, 2007). Consequently, it is neither clear how egg trading can initially spread nor to what extent it can resist invasion by withholders under the high encounter rates that are thought to facilitate its establishment.

Here we build on the analytical model of Henshaw et al. (2014) and extend it by adding four fundamental features. First, we allow for the possible occurrence of withholders, i.e., “cheaters” who never provide eggs and only mate in the male role, in addition to traders and providers. Second, we relax the implicit assumption in Henshaw et al. (2014) that egg production has no costs in terms of availability for mating. This assumption does not generally hold in nature since the time and energy devoted to the acquisition of resources for egg production often trades off with the time and energy available for mate search (Puurtinen & Kaitala, 2002). A direct implication of this trade-off is that individuals who are in the process of producing new eggs are expected to be less available for matings (in the male role since they have no eggs) than individuals carrying eggs. Third, we assume that traders can detect withholders with some positive probability and “punish” them by not providing eggs. Fourth, we incorporate the biologically important feature, discussed by Henshaw et al. (2014) but not incorporated in their model, that eggs might senesce and become unviable before a partner is found. We show that the first three additions generate complex evolutionary dynamics that allow traders to invade (and resist invasion from) both providers and withholders when encounter rates are intermediate and both the costs of egg production and the probability that withholders can be detected are sufficiently high. The fourth addition (egg senescence) shapes the trade-offs that affect the evolution of egg trading.

## Model

We posit a large, well-mixed population of simultaneous hermaphrodites in which generations overlap and there is no self-fertilization. At any given time, each individual in the population is either carrying a batch of eggs or not. Eggless individuals produce a new batch of eggs at a normalized rate of 1. Egg-carrying individuals encounter potential mates at the positive encounter rate *m*, while eggless individuals (who are producing new eggs) encounter potential mates at a discounted rate *λm*, where 0 < *λ* ≤ 1. The parameter *λ* measures the degree to which individuals who are in the process of producing eggs are available for mating. Being unavailable for mating constitutes a cost of egg production in terms of missed opportunities for reproduction in the male role. Thus, low values of mating availability *λ* imply a high cost of egg production, with the extreme case *λ* = 0 implying maximal costs (mating in the male role is impossible while producing eggs). Conversely, high mating availability *λ* implies a low cost of egg production, with *λ* = 1 implying minimal cost (individuals who are in the process of producing new eggs can always mate in the male role). We also incorporate egg senescence, with eggs becoming non-viable at a rate *ρ* ≥ 0.

We consider three different mating strategies: T (“trading”), H (“withholding”), and P (“providing”). All three strategies mate in the male role (i.e., fertilize eggs) whenever possible, but differ on the conditions under which they provide eggs to partners for fertilization. Traders are choosy: they only provide eggs if mates have eggs to reciprocate. Withholders are stingy: they never provide eggs, and only reproduce through their male function. Indeed, the only function of their eggs is to elicit egg release from traders, i.e., withholders “cheat” on their partners by failing to reciprocate eggs. Providers are generous: they provide eggs to any partner, regardless of whether the mate has eggs to reciprocate. We further assume that traders can detect withholders with a positive probability *q*, in which case eggs are not provided. In the absence of withholders (there are only providers and traders in the population) and after setting *λ* = 1 (egg production is costless in terms of availability for mating), and *ρ* = 0 (eggs do not senesce), our model recovers the analytical model of Henshaw et al. (2014), after identifying our “providers” with their “non-traders”.

In line with game-theoretic approaches (Maynard Smith, 1982), we assume a one-locus haploid genetic system, so that each individual’s mating strategy is determined by a single gene inherited from the mother or the father with equal probability. Moreover, we assume a separation of time scales such that the demographic variables (the proportions of individuals carrying and not carrying eggs within each strategy) equilibrate much faster than the evolutionary variables (the proportions of individuals following each strategy). With these assumptions, we can write the evolutionary dynamics of our model as a system of replicator equations (Hofbauer & Sigmund, 1998; Weibull, 1995) for the three strategies T, H, and P, with frequencies respectively given by *x, y*, and *z*. That is, we write the evolutionary dynamics of our model as

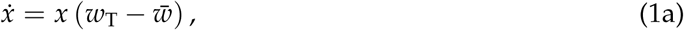

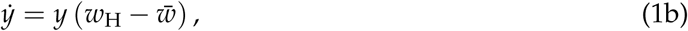

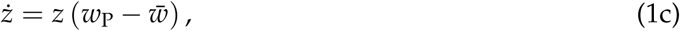

where dots denote time derivatives, *w*_T_, *w*_H_, and *w*_P_ are the fitnesses to each strategy, and 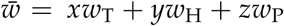 is the average fitness in the population. Fitnesses are given by the rate of offspring production in both the male and the female roles, and are non-trivial functions of the parameters of the model and of the proportions of the different strategies when carrying and not carrying eggs at the demographic equilibrium. The state space Δ is the simplex of all (*x, y, z*) with *x, y, z* ≥ 0 and *x* + *y* + *z* = 1.

In the following we present a summary of our results. Our formal model and the analytical derivation of all results are given in Appendix A: Detailed Model Description and Appendix B: Analysis of the Evolutionary Dynamics.

## Results

The replicator dynamics has three monomorphic equilibria: a homogeneous population of traders (T), a homogeneous population of withholders (H), and a homogeneous population of providers (P). Among these equilibria, H is always unstable: for any parameter combination a homogeneous population of withholders can be invaded by traders, providers, or a mixture of both strategies. In addition to these three monomorphic equilibria, and depending on parameter values, the replicator dynamics can have up to two out of three polymorphic equilibria on the boundary of the simplex Δ (fig. 1): (i) an equilibrium R along the TP-edge, where traders and providers coexist but withholders are absent (figs. 1B, 1C), (ii) an equilibrium Q along the TH-edge, where traders and withholders coexist but there are no providers (figs. 1C, 1D), and (iii) an equilibrium S along the HP-edge, where withholders and providers coexist but where there are no traders (figs. 1D, 1E). When these polymorphic equilibria exist, R is a saddle (repelling for points along the TP-edge, and attracting for neighboring points in the interior of Δ), Q is stable (attracting from neighboring points in Δ), and S is a saddle (attracting for points along the HP-edge, and repelling for neighboring points in the interior of Δ). These equilibria are rather complicated functions of the model parameters, so we report their expressions in Appendix B: Analysis of the Evolutionary Dynamics. The replicator dynamics has no equilibria in the interior of Δ, i.e., no population composition with all three strategies coexisting is an equilibrium.

**Figure 1:**
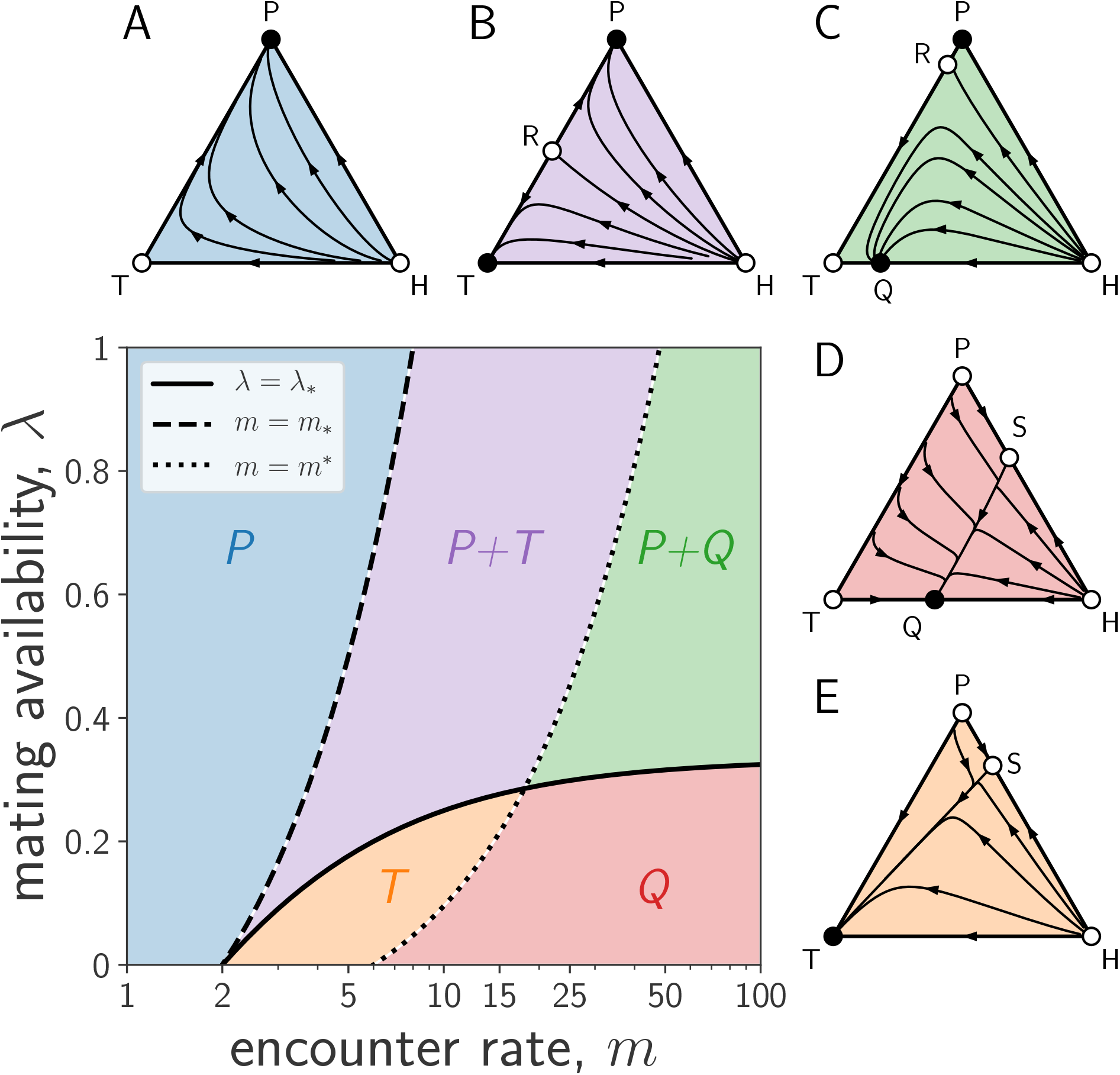
Effects of mating availability and encounter rates on the evolutionary dynamics of egg trading. The parameter space can be divided into five disjoint regions (*P, P+T, P+Q, Q*, and *T*) depending on how availability *λ* compares to the critical availability *λ*_*_ (equation (2)) and on how the encounter rate *m* compares to the critical encounter rates *m*_*_ (equation (3)) and *m*\* (equation (4)). The triangles Δ represent the state space Δ = *{*(*x, y, z*) ≥ 0, *x* + *y* + *z* = 1*}*, where *x, y*, and *z* are the frequencies of traders, withholders, and providers, respectively. The three vertices T, H, and P correspond to homogeneous states where the population is entirely comprised of traders (*x* = 1), withholders (*y* = 1), or providers (*z* = 1). Full circles represent stable equilibria (sinks); empty circles represent unstable equilibria (sources or saddle points). (A) In region *P* trajectories in Δ converge to P. (B) In region *P+T* trajectories converge to either P or T, depending on initial conditions. The equilibrium R on the TP-edge is a saddle point dividing the basins of attraction of P and T. (C) In region *P+Q* trajectories converge to either P or the equilibrium Q along the TH-edge, depending on initial conditions. (D) In region *Q* trajectories converge to Q. The equilibrium S along the HP-edge is a saddle. (E) In region *T* trajectories converge to T. Parameters: *ρ* = 1, *q* = 0.5, *m* = 2 (A), 12 (B), 50 (C and D) or 8 (E), and *λ* = 0.7 (A, B, and C), or 0.1 (D and E).

We find that both the stability of the monomorphic equilibria T and P, and the existence of the polymorphic equilibria Q, R, and S, depend on how the mating availability *λ* compares to the critical value

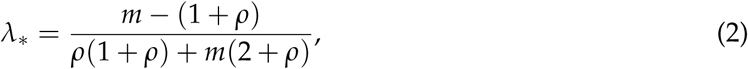

and on how the encounter rate *m* compares to the critical values

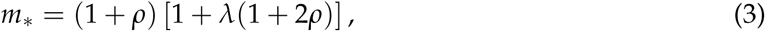

and

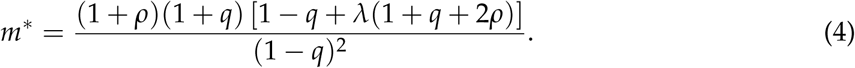

First, the stability of the monomorphic equilibrium P depends on how the mating availability *λ* compares to the critical value *λ*_*_. A homogeneous population of providers is stable against invasions by the other two strategies if and only if mating availability is high (*λ* > *λ*_*_). As *λ* decreases and crosses the threshold *λ*_*_, P becomes unstable against both traders and withholders, and the saddle S is created along the HP-edge.

Second, the stability of the monomorphic equilibrium T depends on how the encounter rate *m* compares to the critical values *m*_*_ and *m*\*. A homogeneous population of traders is: (i) unstable against invasion by providers but stable against invasion by withholders if the encounter rate is low (*m* < *m*_*_), (ii) stable against both withholders and providers if the encounter rate is intermediate (*m*_*_ < *m* < *m*\*), and (iii) stable against invasion by providers but unstable against invasion by withholders if the encounter rate is high (*m* > *m*\*). As *m* increases and crosses the threshold *m*_*_, T becomes stable while spawning the unstable equilibrium R along the TP-edge; as *m* increases further and crosses the threshold *m*\*, T becomes unstable and the stable equilibrium Q (where traders and withholders coexist) is created along the TH-edge.

All in all, the parameter space can be partitioned into five dynamical regions (fig. 1), each having qualitatively different evolutionary dynamics. Among these, only regions *Q* and *T* (for which availability is low, i.e., *λ* < *λ*\* holds) allow traders to invade a resident population of providers, and only region *T* allows traders to both invade providers and resist invasion by withholders. A key requirement for this last scenario is that encounter rates are neither too high nor too low (*m*_*_ < *m* < *m*\*).

The encounter rate *m* is a key parameter in our model. For low encounter rates (*m* < *m*_*_; region *P*), P is the only stable equilibrium and the outcome of the evolutionary dynamics. This makes intuitive sense: if potential mates are difficult to find, individuals should provide eggs at every mating opportunity; being picky in this context is risky as another partner might be difficult to find before eggs become unviable. For higher encounter rates (*m* > *m*_*_; regions *P+T, T, P+Q*, and *Q*) finding mates becomes easier, and it pays to reject eggless partners in the hope of finding partners carrying eggs. Very large encounter rates (*m* > *m*\*; regions *P+Q* and *Q*) even allow withholders (who never release their eggs and only mate in the male role) to be successful in the long run and coexist with traders at the equilibrium Q. The proportion of traders at such an equilibrium decreases as the mate encounter rate increases, down to 50% in the limit of high encounter rates.

The benefits of being choosy are particularly salient when the costs of egg production are high (i.e., when the mating availability *λ* is low). Indeed, a lower mating availability *λ* has two related and reinforcing consequences. First, low availability means fewer opportunities to mate in the male role when not carrying eggs, and hence higher opportunity costs to mate indiscriminately in the female role. Second, low availability also implies that the probability of finding another potential mate without eggs after having rejected previous potential partners is lower, thus decreasing the risk of being choosy. In line with these arguments, we find that for sufficiently high costs of egg production (*λ* < *λ*_*_; regions *Q* and *T*), P can be invaded by strategies that do not mate indiscriminately in the female role (traders and withholders). For high encounter rates (*m* > *m*\*; region *Q*) traders invade but are not able to displace withholders, and the population composition at equilibrium is a mixture of traders and withholders. Otherwise, for moderate encounter rates (*m*_*_ < *m* < *m*\*; region *T*) traders invade and take over the whole population while resisting invasion by withholders.

The probability that traders detect withholders, *q*, plays an essential role in stabilizing the trading equilibrium T in our model (fig. 2). Indeed, some amount of withholder detection (as encapsulated by the parameter *q*) is necessary for trading to be evolutionarily stable in the presence of withholders. This is so because the critical encounter rate *m*\* tends to *m*_*_ (which does not depend on *q*) as *q* tends to zero. Thus, in this limit, regions *P+T* and *T* cease to exist and the trading equilibrium T is unstable for all encounter rates. In addition, the critical encounter rate *m*\* is an increasing function of *q* (fig. 2). As *m* ≤ *m*\* is a necessary and sufficient condition for a monomorphic population of traders to resist invasion by withholders, larger values of *q* imply that more stringent conditions (i.e., higher encounter rates) are required to destabilize T.

**Figure 2:**
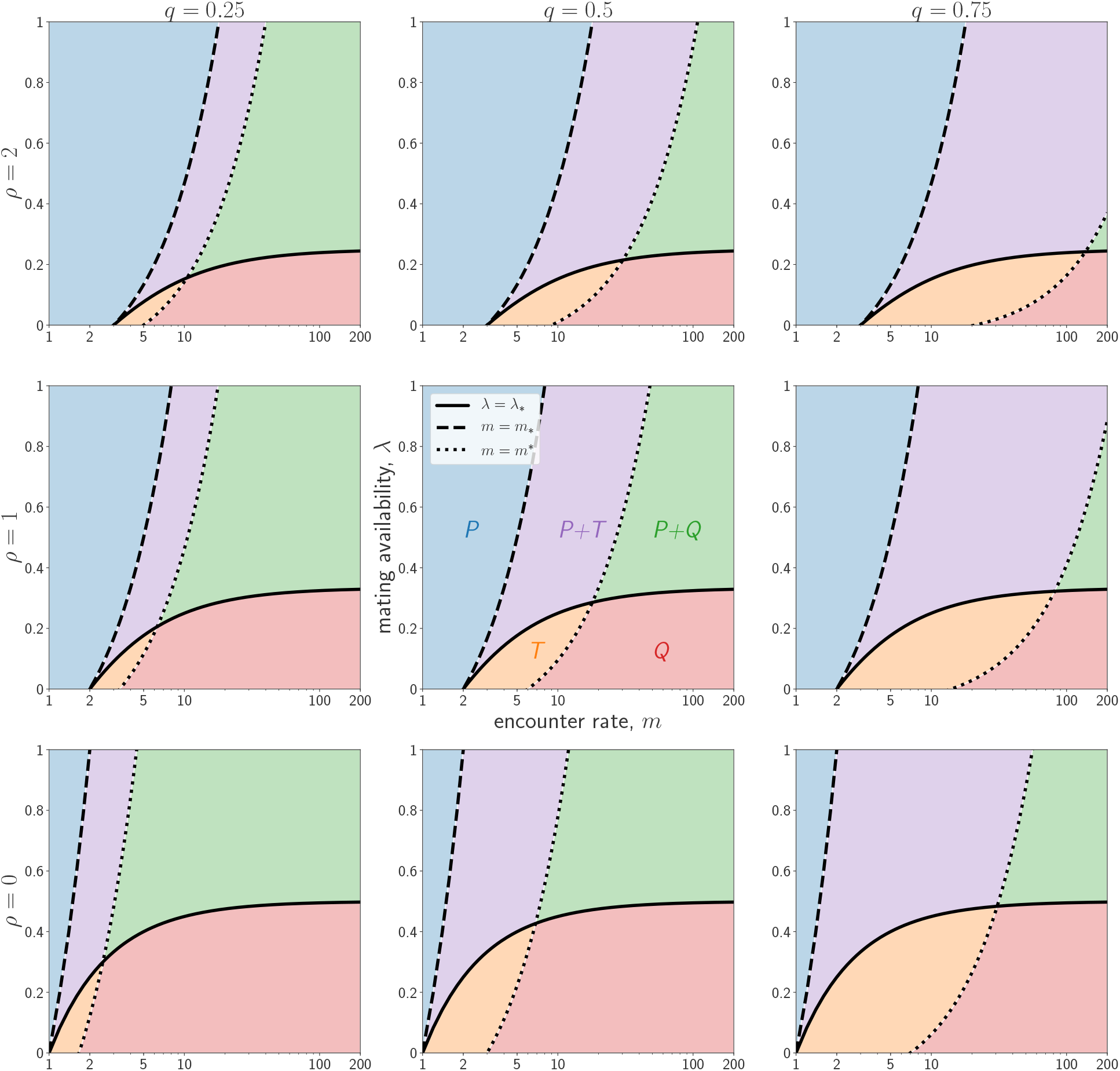
Effects of egg senescence and probability of withholder detection on the evolutionary dynamics of egg trading. Panels represent, for different combinations of egg senescence *ρ* and probability of withholder detection *q*, the critical mating availability *λ*_*_ (equation (2)), and the critical encounter rates *m*_*_ (equation (3)) and *m*\* (equation (4)) that define the boundaries of the five dynamical regions (*P, P+T, P+Q, Q*, and *T*) into which the parameter space can be divided. For fixed *ρ* and *λ*, increasing *q* increases the values of the encounter rate *m* at which *m* = *m*\* holds, thus increasing the areas of regions *P+T* and *T* (where the trading equilibrium T is evolutionarily stable) and shrinking the areas of regions *P+Q* and *Q* (where withholders invade T). For fixed *q* and *m*, increasing *ρ* decreases the values of the mating availability *λ* at which *λ* = *λ*_*_ holds, thus decreasing the combined area of regions *Q* and *T*, where traders can invade the providing equilibrium P. The middle panel (second row, second column) corresponds to the parameter values (*ρ* = 1, *q* = 0.5) used in fig. 1.

Finally, we note that the critical mating availability *λ*_*_ and the critical encounter rates *m*_*_ and *m*\* are all functions of the rate of egg senescence *ρ*. The critical availability *λ*_*_ is decreasing in *ρ* (fig. 2). The evolutionary consequence of this effect is that the higher the rate of egg senescence *ρ*, the lower the critical availability *λ*_*_ below which traders (and withholders) can invade a monomorphic population of providers. This makes intuitive sense as providers give up their eggs more freely and are thus less likely to suffer the consequences of a higher egg senescence than traders and withholders. Additionally, both critical encounter rates *m*_*_ and *m*\*are increasing in *ρ* (fig. 2). Therefore, the higher *ρ*, the higher the minimal encounter rate *m*_*_(respectively, the maximal encounter rate *m*\*) required for a monomorphic population of traders to resist invasion by providers (respectively, by withholders).

## Discussion

A general prediction of our model is that there are only three possible evolutionarily stable equilibria: a homogeneous population of providers, a homogeneous population of egg traders, or a polymorphic population that includes both egg traders and withholders. The first stable equilibrium would correspond to simultaneous hermaphrodites that do not trade eggs. This equilibrium is attained in a large area of the parameter space, which is consistent with the fact that the majority of simultaneous hermaphrodites do not trade eggs. The second stable equilibrium would correspond to egg traders and can be attained under the specific conditions that we discuss below. The closest situation to the third stable equilibrium in nature would correspond to egg-trading species in which mating also occurs in the male role only through streaking, i.e., the furtive release of sperm in competition with the male of an egg trading pair (Fischer, 1984; Oliver, 1997; Petersen, 1995; Pressley, 1981). Streaking was not explicitly incorporated in our model but we note that, as our withholders, such streakers are not pure males but simultaneous hermaphrodites that mate in the male role. We are not aware of simultaneously hermaphroditic species in which egg trading is facultative, which is consistent with the fact that there is no stable equilibrium in our model involving both traders and providers.

When mating availability (*λ*) is equal to one and egg senescence (*ρ*) is equal to zero, the only difference between our model and the one in Henshaw et al. (2014) is that we incorporate withholders. Doing so does not affect the conclusion from Henshaw et al. (2014) that there is an initial barrier that traders need to overcome in order to invade a population of providers. Further, as predicted by Henshaw et al. (2014), higher encounter rates make this invasion barrier smaller and in this sense high encounter rates thus promote the evolution of egg trading. However, very high encounter rates (*m* > *m*\*) will also inevitably allow withholders to invade the trading equilibrium and thereby lead to the emergence of a stable mixture of traders and withholders. In particular, in the limit of very high encounter rates (so that the invasion barrier becomes arbitrarily small) the evolutionary outcome is not the invasion and fixation of trading predicted by Henshaw et al. (2014), but (as we show in Appendix B: Analysis of the Evolutionary Dynamics) a stable mix consisting of 50% traders and 50% withholders. Such a mix is stable because with very high encounter rates withholders prosper in a population where there are ample opportunities to reproduce in the male role (as will be the case if traders, who are willing to provide their eggs with probability 1–*q*, are frequent) while they fare poorly in a population with few opportunities to reproduce in the male role (as will be the case in a population consisting predominantly of withholders who never provide their eggs).

Recognizing the possibility of costly egg production by allowing mating availability to be less than one is another important way in which our model differs from Henshaw et al. (2014). Indeed, our analysis reveals that the cost of egg production plays a crucial role in the evolution of egg trading. In particular, for encounter rates that are neither too high nor too low, traders can both (i) invade providers at sufficiently high encounter rates, and (ii) be stable against invasion by withholders at sufficiently small encounter rates. This result implies that neither a combination of self-fertilization and kin selection (Axelrod & Hamilton, 1981) nor high encounter rates (Henshaw et al., 2014) that would promote the invasion by withholders are necessary for the evolution of egg trading, and thereby resolves the dilemma on the relationship between encounter rate and the evolution of egg trading.

The trade-off between the time and energy allocated to acquire resources for egg production versus mate search that is captured by our parameter *λ* has been documented in egg traders. For example, in the hamlets (*Hypoplectrus* spp.), one of the fish groups in which egg trading is best described, individuals meet on a daily basis in a specific area of the reef for spawning at dusk (Fischer, 1980). This can imply swimming over hundreds of meters of reef (Puebla et al., 2012). Not all individuals show up in the spawning area on each evening, but most individuals that are present are observed spawning in both the female and male role (implying that they carry eggs). The majority of individuals who do not spawn are not present in the spawning area and are therefore not available for mating, even in the male role only, which is exactly what the parameter *λ* captures. This said, our model is not meant to represent any group of egg traders in particular but to capture the minimal set of parameters that are relevant for the evolution of egg trading. Mate encounter rate had been identified as such a parameter by Henshaw et al. (2014); we added here the opportunity costs of egg production. Our results indicate that the evolution of egg trading from an ancestral state where the population consists only of providers requires at the very least a minimum of egg-production costs.

Once egg trading is able to invade a population of providers, two different evolutionary scenarios are possible. First, trading can reach fixation and be established at an evolutionarily stable equilibrium. Second, trading can be sustained at a polymorphic equilibrium featuring egg traders and withholders. Which of these two scenarios is reached depends to a large extent on the ability of egg traders to detect withholders (*q*). A necessary condition for the first scenario to be reached is that *q* is positive, i.e., that there is at least some withholder detection. Moreover, the higher *q* (i.e., the better the abilities of traders to detect withholders), the larger the set of values for the other parameters under which trading is evolutionarily stable against withholding and the first scenario prevails.

There are at least two ways in which egg traders may be able to detect withholders in nature. The first one is through reputation and learning in small populations where mating encounters occur repeatedly among the same set of individuals (Puebla et al., 2012). In this situation, individuals who fail to reciprocate eggs might be identified as withholders and avoided in subsequent mating encounters. The second one is through parcelling of the egg clutch, which occurs in several egg-trading species (Fischer, 1980; Fischer & Hardison, 1987; Oliver, 1997; Petersen, 1995). In this case eggs are divided into parcels that the two partners take turns in providing and fertilizing. This constitutes an efficient mechanism to detect partners that fail to reciprocate, and also provides the opportunity to terminate the interaction before all eggs are released if the partner does not reciprocate.

By and large, the conditions that are required for the invasion and fixation of egg trading (intermediate encounter rates, sufficiently high costs of egg production and possibility to detect withholders) are rather restrictive. In addition, egg trading requires that individuals interact directly to trade eggs, which implies that they are mobile. It is therefore not surprising that egg trading is a rare mating system, documented only in Serraninae fishes (Fischer, 1980, 1984; Oliver, 1997; Petersen, 1995; Pressley, 1981) and dorvilleid polychaetes in the genus *Ophryotrocha* (Sella, 1985; Sella & Lorenzi, 2000; Sella et al., 1997; Sella & Ramella, 1999). Hermaphroditism, on the other hand, occurs in 24 out of 34 animal phyla and is common to dominant in 14 phyla including sponges, corals, jellyfishes, flatworms, mollusks, ascidians and annelids (Jarne & Auld, 2006). The rare occurrence of egg trading among simultaneous hermaphrodites suggests that simultaneous hermaphroditism can readily evolve and be maintained in the absence of egg trading. This is what motivated our choice to focus on the evolution of egg trading among simultaneous hermaphrodites as opposed to the joint evolution of egg trading and simultaneous hermaphroditism. In our model this is illustrated by the fact that although withholders mate in the male role exclusively, they are nonetheless not pure males but hermaphrodites that keep producing eggs to elicit egg release by traders. In principle, the rarity of egg trading might also be due to the possibility that egg trading ultimately leads to a loss of hermaphroditism and consequently of egg trading itself. However, this scenario goes against the results of Henshaw et al. (2015), who show that egg trading can help stabilizing hermaphroditism by selecting for a female-biased sex allocation in traders, which in turn prevents pure females from invading a population of traders.

We assumed a very simple genetic architecture of the trait under consideration, namely a one-locus haploid genetic system. Since most simultaneously hermaphroditic species are diploid, and since egg trading is likely to be a complex trait under the control of many genes, this is clearly a simplifying assumption that trades biological reality for model tractability, i.e., an example of the “phenotypic gambit” often endorsed in evolutionary models (Gardner et al., 2011; Grafen, 1984). In our case, this simplifying assumption is justified both by the fact that the specific genetic architecture of egg trading is so far unknown for any species, and by our goal of comparing our model and results with the existing literature, which has also explicitly or implicitly endorsed the phenotypic gambit. This said, egg trading and other traits affecting mating strategies are particular because they influence who mates with whom and can thus potentially lead to assortment of alleles at the zygotic level. Additional work is needed to investigate the effect of the genetic system (e.g., number of loci, dominance) on the evolutionary dynamics of egg trading.

A key dynamic that is characteristic of systems subject to sexual conflict over mating such as the one investigated here is the co-evolution of male coercion and female resistance (Clutton-Brock & Parker, 1995). While male coercion has been considered in the context of egg trading (Fischer & Hardison, 1987), there is little evidence of this phenomenon among egg traders. Nevertheless, the streaking behavior displayed by some egg trading species (Fischer, 1984; Oliver, 1997; Petersen, 1995; Pressley, 1981) may be interpreted as a form of male coercion. Henshaw et al. (2014) included streaking in their simulation model and found that it makes the evolution of egg trading less likely (see also Henshaw et al. (2015) for the effects of streaking on the role played by egg trading in stabilizing hermaphroditism). This is because streakers (as our withholders) bypass the trading convention and gain reproductive success as males without offering eggs in return. This form of male coercion could be counteracted by strategies of female resistance that increase the costs of coercion, such as the parcelling of the egg clutch observed in some egg traders (Fischer, 1980; Fischer & Hardison, 1987; Oliver, 1997; Petersen, 1995). This calls for the incorporation of both streaking and egg parcelling in future analytical models to better understand the evolution of egg trading.

We modeled social interactions as a game with three distinct strategies (traders, providers, and withholders) and analyzed the resulting evolutionary process using the replicator dynamics. A caveat of this approach is that withholding is an evolutionary dead end, as a population of withholders would completely fail to reproduce. This may cast some doubt on the suitability of modelling withholding as a pure strategy, and on the results we obtained. To dispel this potential criticism and to test the robustness of our results, Appendix C: Alternative Model with Probabilistic Withholding presents a model where traders compete against non-traders playing a mixed strategy that provides eggs with probability *s* and withholds them with probability 1 − *s*. We use adaptive dynamics (Doebeli, 2011; Geritz et al., 1998) to determine the evolutionary end-point of the quantitative trait *s* in a population of non-traders, and then investigate the conditions under which traders are able to invade such a population. The results of this analysis demonstrate the robustness of our conclusion that traders can invade non-trading populations if the encounter rate is intermediate and mating availability is sufficiently low. A more ambitious analysis could fully embrace a continuous representation of the phenotype space and use multidimensional versions of adaptive dynamics (e.g., Débarre et al. (2014); Leimar (2009); Mullon et al. (2016)) to investigate the coevolution of rates of providing, withholding, and trading eggs in a relatively economic way.

Our model of egg trading is related to game-theoretic models of food sharing and social foraging where individuals either “produce” by searching for food or “scrounge” by not searching and instead exploiting others’ food discoveries (Giraldeau & Caraco, 2000; Vickery et al., 1991). In these producer-scrounger games, it is assumed (i) that information about the location of food clumps discovered by producers is immediately acquired by scroungers, and (ii) that single individuals are able to process the food clumps they discover. Our model of egg trading can be thought of as a variant of a producer-scrounger game in which information about the location of resources is instead private, where two individuals are needed to access or handle a resource (e.g., large prey), and where “mating” corresponds to entering a partnership to successfully exploit the resource. From this perspective, the providers of our model are equivalent to producers that search for food and share information with all individuals; withholders to scroungers that either do not search or always withhold information; and traders to individuals that search but only share information on discovered food items with partners that have acquired new information.

Our model of egg trading also shares features with more general models for the evolution of cooperation, in particular with models of partner choice (Bull & Rice, 1991; Noë & Hammerstein, 1994) and indirect reciprocity (Nowak & Sigmund, 2005). First, our model is related to models of partner choice where potential partners are encountered at a certain rate, and where strategies or individuals can vary both in their “choosiness” and in their “cooperativeness” (e.g., André & Baumard (2011); McNamara et al. (2008)). Importantly, however, in our model individuals discriminate partners not directly on the basis of their perceived cooperativeness, but rather on their state or physiological condition (i.e., on whether or not the partner is carrying eggs), which serves as an indirect measure of partner quality. Second, the transitions a given focal individual makes between different states or physiological conditions are mediated by social actions, e.g., an egg carrier becomes eggless when it decides to offer its eggs to a partner. This resembles the way models of indirect reciprocity work, where an individual’s reputation changes depending on both its decision to cooperate and the particular social norms to assign reputations enforced in the population (e.g., Leimar & Hammerstein (2001); Nowak & Sigmund (1998); Ohtsuki & Iwasa (2006); Panchanathan & Boyd (2003); Santos et al. (2018)).

Our model predicts that egg trading should occur in simultaneously hermaphroditic species for which encounter rates are intermediate, egg production entails a cost in terms of mating availability, and withholders can be detected to some extent. Testing this prediction calls for an empirical estimation of these factors (as well as rates of egg senescence) in egg-trading and closely related non-egg-trading species. The incorporation of egg parcelling and sperm competition through streaking into our model would also allow to refine our predictions.

## Acknowledgments

We thank Erol Akçay and two anonymous reviewers for comments that helped to improve this manuscript. This research was funded by a Future Ocean Cluster of Excellence grant to O. Puebla. J. Peña acknowledges IAST funding from the French National Research Agency (ANR) under the Investments for the Future (Investissements d’Avenir) program, grant ANR-17-EURE-0010. The code used for creating the figures of this paper builds on Inom Mirzaev and Drew F. K. Williamson’s Python package egtplot (https://github.com/mirzaevinom/egtplot). Our source code in Python is publicly available on GitHub (https://github.com/jorgeapenas/eggtrading).

## Appendix A: Detailed Model Description

Our model builds on the analytical model of Henshaw et al. (2014), extending it in a number of directions.

We posit a large, well-mixed population of simultaneous hermaphrodites. At any time, each individual in the population either is or is not carrying a batch of eggs. Individuals without eggs produce a new batch of eggs at a rate normalized to 1, so that all other rates are measured relative to the rate of egg production. Potential mates are encountered at rate *m* > 0 if the focal individual carries eggs, or at a discounted rate *λm*, where 0 < *λ* ≤ 1, if the focal does not carry eggs. Equivalently, an individual not carrying eggs is available for encounters with probability *λ*. Hence, *λ* captures the opportunity costs of egg production; *λ* < 1 means that an individual busy producing eggs cannot be available all the time as a potential partner in the male role. We assume that eggs senesce and become unviable at rate *ρ* ≥ 0.

Individuals adopt one of three possible strategies: trading (T), withholding (H), or providing (P). Our traders behave like the traders in Henshaw et al. (2014): they offer their eggs only to partners carrying eggs (who can reciprocate). Withholders produce and carry eggs but never release them to partners, thereby only reproducing through the male role. Providers correspond to the “non-traders” in Henshaw et al. (2014): they offer their eggs to any partner (either carrying or not carrying eggs). All three strategies fertilize the eggs offered to them by partners. Finally, we assume that traders can detect withholders with probability 0 < *q* < 1 and “punish” them by not releasing eggs.

Normalizing the value of a fertilized batch of eggs to 1, table A1 summarizes the resulting “payoffs” (i.e., the reproductive success arising from mating in the female and the male role) of each strategy in each individual state (i.e., carrying or not carrying a batch of eggs). For instance, when a trader carrying eggs meets a withholder carrying eggs, then the trader will agree to mate with probability 1 *-q* and in this case have its batch of eggs fertilized by the withholder, while the batch of eggs of the withholder is not released. As a result, the trader gets a payoff of 1 – *q* through the female role and a payoff of 0 through the male role (first row, second column of table A1), while the withholder gets a payoff of 0 through the female role and a payoff of 1 *-q* through the male role (second row, first column of table A 1). Other payoff values are calculated in a similar way.

The model in Henshaw et al. (2014) is recovered from our general model by (i) allowing only for providers and traders, (ii) assuming costs of egg production are zero (by setting *λ* = 1), and (iii) ignoring egg senescence (by setting *ρ* = 0).

### Proportions of strategies and of egg carriers

Let *x, y*, and *z* denote the respective proportions of traders, withholders, and providers in the population, satisfying

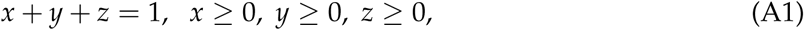

and let Δ denote the set of population shares (*x, y, z*) of the three strategies satisfying the conditions in (A1). Similarly, let *x*_*e*_, *y*_*e*_, and *z*_*e*_ denote the proportions (relative to the overall population size) of, respectively, traders carrying eggs, withholders carrying eggs, and providers carrying eggs, with the corresponding proportions of individuals not carrying eggs given by

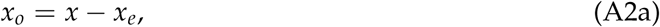

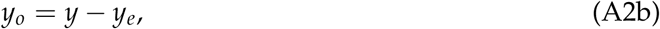

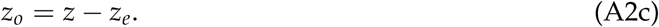

To abbreviate formulas, it will sometimes be convenient to use *e* and *o* to denote the population fractions carrying eggs, resp. not carrying eggs:

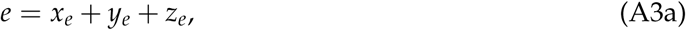

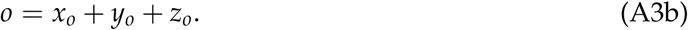

**Table A1:** Reproductive success (or “payoffs”) to a given focal individual (rows) when encountering a given partner (columns). The first entry in each cell of the matrix corresponds to reproduction in the female role, while the second entry corresponds to reproduction in the male role. Payoffs are normalized so that the value of one batch of eggs is equal to 1. T_*e*_ (resp. T_*o*_) indicates a trader carrying eggs (resp. not carrying eggs). A similar convention applies to the other two strategies, with H_*e*_ and H_*o*_ indicating withholders carrying and not carrying eggs, respectively, and P_*e*_ and P_*o*_ indicating providers carrying and not carrying eggs, respectively.

|  |  | Partner |  |  |  |  |  |
| --- | --- | --- | --- | --- | --- | --- | --- |
| | | $T_e$ | $H_e$ | $P_e$ | $T_o$ | $H_o$ | $P_o$ |
| Focal | $T_e$ | 1,1 | $1 - q, 0$ | 1,1 | 0,0 | 0,0 | 0,0 |
| | $H_e$ | $0, 1 - q$ | 0,0 | 0,1 | 0,0 | 0,0 | 0,0 |
| | $P_e$ | 1,1 | 1,0 | 1,1 | 1,0 | 1,0 | 1,0 |
| | $T_o$ | 0,0 | 0,0 | 0,1 | 0,0 | 0,0 | 0,0 |
| | $H_o$ | 0,0 | 0,0 | 0,1 | 0,0 | 0,0 | 0,0 |
| | $P_o$ | 0,0 | 0,0 | 0,1 | 0,0 | 0,0 | 0,0 |

**Table A2:** Encounter rates of potential partners for a focal individual that is carrying eggs (*e*, first row) or not carrying eggs (*o*, second row).

|  |  | Partner |  |  |  |  |  |
| --- | --- | --- | --- | --- | --- | --- | --- |
| | | $T_e$ | $H_e$ | $P_e$ | $T_o$ | $H_o$ | $P_o$ |
| Focal | $e$ | $m \frac{x_e}{e+\lambda o}$ | $m \frac{y_e}{e+\lambda o}$ | $m \frac{z_e}{e+\lambda o}$ | $m \frac{\lambda x_o}{e+\lambda o}$ | $m \frac{\lambda y_o}{e+\lambda o}$ | $m \frac{\lambda z_o}{e+\lambda o}$ |
| | $o$ | $\lambda m \frac{x_e}{e+\lambda o}$ | $\lambda m \frac{y_e}{e+\lambda o}$ | $\lambda m \frac{z_e}{e+\lambda o}$ | $\lambda m \frac{\lambda x_o}{e+\lambda o}$ | $\lambda m \frac{\lambda y_o}{e+\lambda o}$ | $\lambda m \frac{\lambda z_o}{e+\lambda o}$ |

### Encounter rates

The rate at which a focal individual encounters potential mates has been assumed to depend on whether the focal is carrying eggs (encounter rate *m*) or not (encounter rate *λm*). As a fraction *e* + *λo* of all individuals are available for meetings, a fraction *x*_*e*_/(*e* + *λo*) of these meetings will be with a trader carrying eggs, resulting in the encounter rates of a focal with a trader carrying eggs given in the first column of table A2. The other encounter rates, summarized in table A2, are obtained by analogous reasoning: in each case the encounter rate of the focal is multiplied by the probability that a partner who is available for a meeting follows the specified strategy and is in the specified state (carrying or not carrying eggs).

### Female and male reproductive success

Female and male reproductive success of the different strategies (traders, withholders, providers) in the different states (carrying or not carrying eggs) can be calculated by combining the information given in tables A1 and A2, as specified in the following.

#### Traders

Consider a focal individual that is a trader and carries a batch of eggs. The expected reproductive success of such an individual through the female function, that we denote by 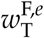, is obtained by multiplying (i) the rate at which a trader carrying eggs encounters each type of potential partner (first row of table A2) by (ii) the expected number of its egg batches such a partner fertilizes (first entry in the corresponding column of the first row of table A1), and summing up all contributions. This yields

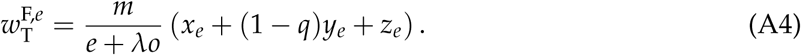

Likewise, the expected reproductive success of a trader carrying eggs through the male function, that we denote by 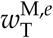, is obtained by multiplying the rate at which a trader carrying eggs encounters each type of potential partner (first row of table A2) by the expected number of the potential partner’s egg batches it gets to fertilize (second entry in the corresponding column of the first row of table A1), and summing up all contributions. This yields

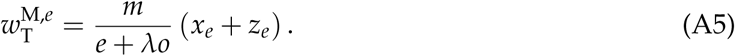

If the focal trader is not carrying eggs, its reproductive success through the female function is obviously

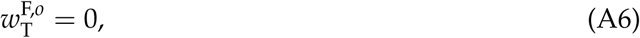

while its reproductive success through the male function is given by

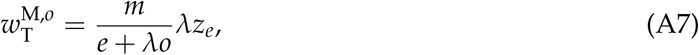

where, similarly to the derivation of (A5), we have combined the information given in the second row of table A2 with that of the fourth row of table A1 to obtain (A7).

#### Withholders

Following a similar procedure as in the case of traders, and employing a similar notation, we obtain the following expressions for the reproductive success of a focal withholder through female and male functions, when carrying or not carrying eggs:

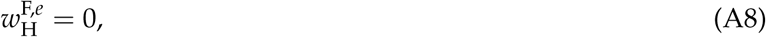

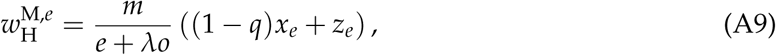

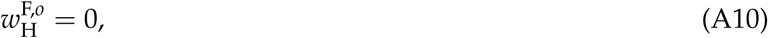

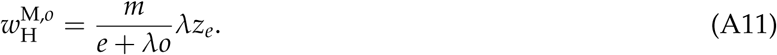

In particular, note that as withholders never release eggs, they gain no reproductive success through the female function.

#### Providers

Similarly, for providers we obtain

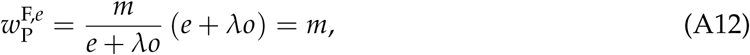

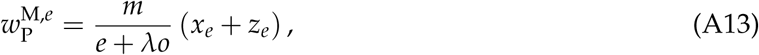

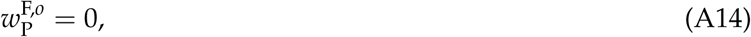

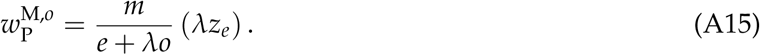

### Equality of female and male reproductive values

In our model, as in models with two separate sexes, all offspring have one mother and one father, so that at every instance the total number of offspring fathered in the male role must equal the total number of offspring mothered in the female role (Fisher, 1930; Fromhage et al., 2016; Houston & McNamara, 2002). With our notation, this means that

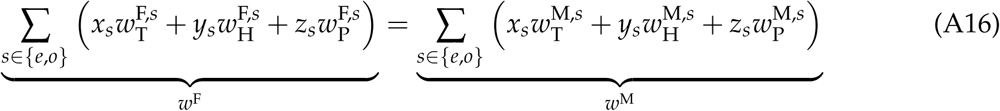

must hold, where *w*^F^ is the population average of the rate of reproduction in the female role and *w*^M^ is the population average of the rate of reproduction in the male role. To verify that this fundamental consistency condition holds, we can substitute from (A4) – (A15) and simplify to obtain

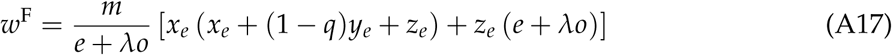

and

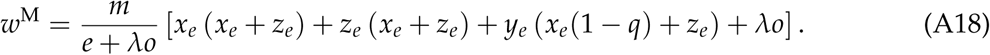

Straightforward algebra establishes that these two expressions are identical, so that condition (A16) holds.

### Demographic dynamics and equilibrium

From our assumptions on the encounter rates of different individuals in the population (table A2), we obtain the following system of differential equations adjusting the proportions of egg-carriers within each subpopulation of strategists:

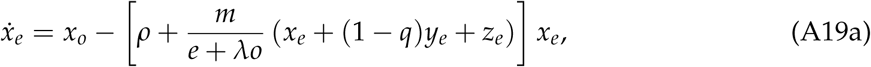

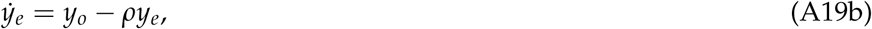

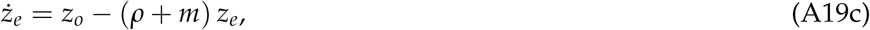

where dots denote time derivatives. The first (positive) term on the right hand side of these equations gives the rate of flow into the egg-carrying state, and is equal to the proportion of individuals not carrying eggs following the relevant strategy (times the rate at which individuals produce eggs, that we have normalized to 1). The second (negative) term on the right hand side gives the outflow from the egg-carrying state. Individuals lose their eggs either because of senescence (explaining the terms proportional to *ρ*) or because they offer their eggs to partners (explaining the terms proportional to *m*). As withholders never give up their eggs when meeting a partner, they only lose eggs due to senescence, so that the outflow from the egg-carrying state is simply *ρy*_*e*_. Egg-carrying providers lose their eggs at rate *m* due to encountering partners, as each encountered partner accepts (i.e., fertilizes) the eggs offered by a provider. This explains the form of (A19c). To understand the outflow from the egg-carrying state for traders due to offering up eggs for fertilization, observe that in a meeting with another individual an egg-carrying trader only gives up its eggs if its partner is also carrying eggs and is not identified as a withholder. Hence, the proportion of meetings in which an egg-carrying trader provides eggs is given by the proportion of meetings in which this condition is satisfied. As a fraction *e* + *λo* of the individuals in the population are available for meetings, this proportion is given by (*x*_*e*_ + (1 − *q*)*y*_*e*_ + *z*_*e*_)/(*e* + *λo*).

Setting the right hand side of (A19) to zero we find that the demographic equilibrium satisfies

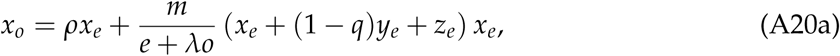

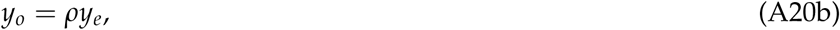

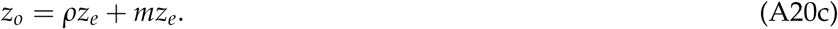

Substituting from (A2) and (A3a), we can rewrite the steady-state equations (A20) solely in terms of (*x, y, z*) and (*x*_*e*_, *y*_*e*_, *z*_*e*_) as

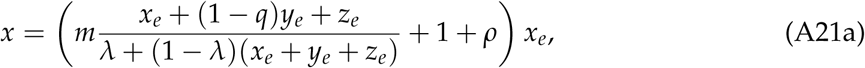

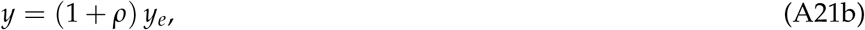

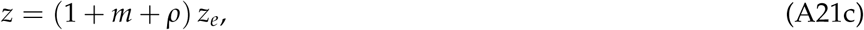

For any (*x, y, z*) ∈ Δ the equations in (A21) have a unique non-negative solution (*x*_*e*_, *y*_*e*_, *z*_*e*_) given by

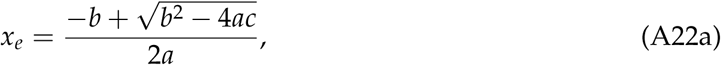

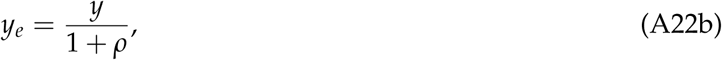

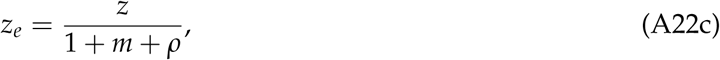

where

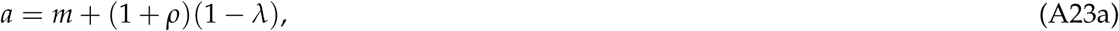

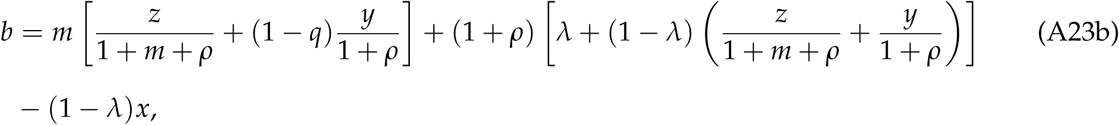

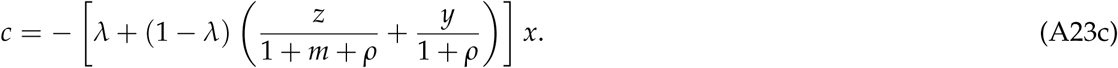

Equations (A22b) and (A22c) are immediate from (A21b) and (A21c). To obtain (A22a), we rewrite (A20a) as

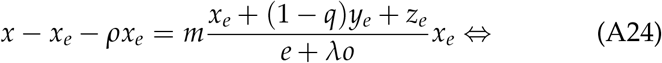

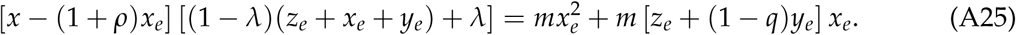

Rearranging, substituting for *y*_*e*_ and *z*_*e*_ from (A22b) and (A22c), and using the definitions in (A23), we obtain that *x*_*e*_ is given by the unique non-negative solution of the quadratic equation

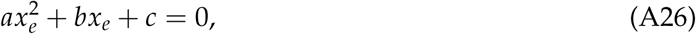

i.e., *x*_*e*_ is given by (A22a).

Before proceeding, we note that (A21) can be rearranged as

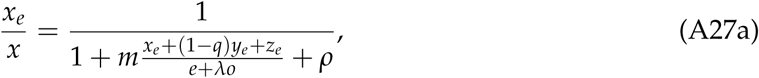

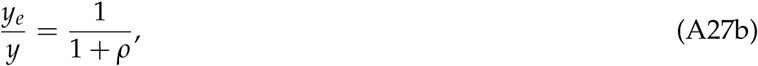

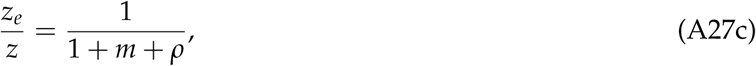

whenever the population share of the strategy under consideration is strictly positive. This gives us expressions for the fraction of time that an individual following one of these strategies is carrying eggs or, alternatively, for the probability that a randomly chosen individual of a given strategy is carrying eggs. When the population share of a strategy is zero, we interpret the expressions *x*_*e*_/*x, y*_*e*_/*y*, and *z*_*e*_/*z* as the corresponding limits (which are well-defined) of the expressions on the right side (A27) as the population share of a strategy goes to zero.

### Expected total reproductive success for each strategy at the demographic equilibrium

We assume a separation of time scales such that the demographic dynamics adjusting the proportions of egg-carriers to their equilibrium values (uniquely determined by (A21)) are much faster than the evolutionary dynamics adjusting the proportions of different strategies in the population. For the evolutionary dynamics we take the rate of offspring produced via reproduction in both sex roles as our fitness measure. Hence, for given frequencies *x, y*, and *z*, we take the fitness of each strategy to be the expected total reproductive success at the demographic equilibrium values *x*_*e*_, *y*_*e*_, and *z*_*e*_, given by (A22). These fitnesses are calculated as follows.

#### Traders

At the demographic equilibrium, a focal trader will be carrying eggs with probability *x*_*e*_/*x* and not carrying eggs with probability *x*_*o*_/*x*. The total expected reproductive success of a trader (through both the female and the male functions) is then given by

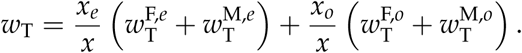

Substituting from (A4)-(A7) and simplifying, we obtain

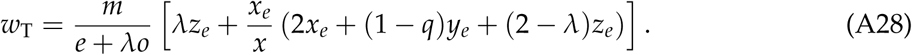

#### Withholders

At the demographic equilibrium, a focal withholder will be carrying eggs with probability *y*_*e*_/*y* and not carrying eggs with probability *y*_*o*_/*y*. The total expected reproductive success of a with-holder is then

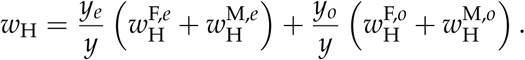

Substituting from (A8)-(A11) and simplifying, we obtain

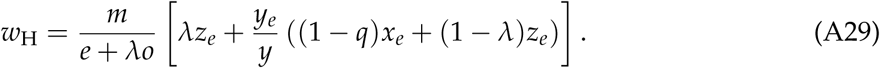

#### Providers

For a focal provider, we obtain

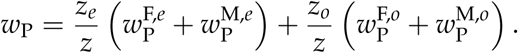

Substituting from (A12)-(A15) and simplifying, we obtain

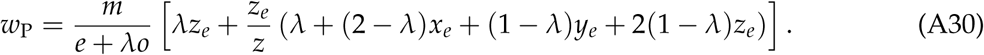

### Evolutionary dynamics

To model the evolutionary dynamics, we make use of the replicator dynamics (Hofbauer & Sigmund, 1998; Weibull, 1995) with total (expected) fitness in the place of expected payoffs. That is, we consider the following system of differential equations:

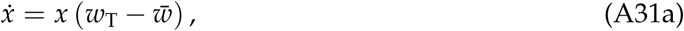

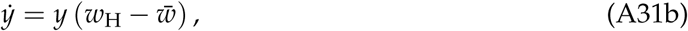

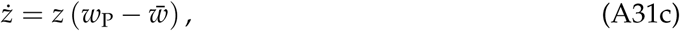

where dots denote time derivatives and

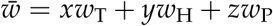

is the average fitness in the population. The frequencies *x*, *y*, *z* can vary within the simplex Δ defined by (A1).

The replicator dynamics is invariant to transformations that either add the same function to all payoffs or multiply payoffs by the same positive function (this last invariance up to a change of speed). We can then subtract the common term *mλz*_*e*_/(*e* + *λo*) from the expressions for *w*_T_, *w*_H_, and *w*_P_ given in equations (A28), (A29), and (A30), and then multiply the resulting expressions by (*e* + *λo*)/*m* to obtain the renormalized fitnesses (which, with slight abuse of notation, we continue to denote by *w*_P_, *w*_T_, and *w*_H_):

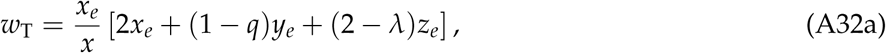

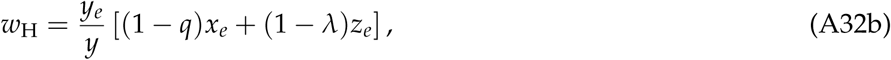

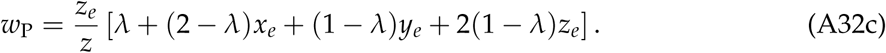

Introducing the abbreviations

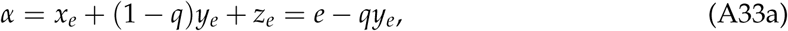

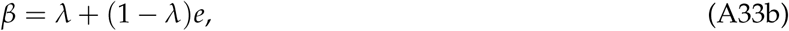

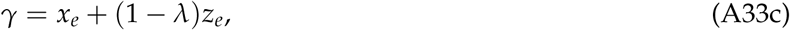

where the second equality in the first line follows from the definitions in (A3), we can rewrite (A32) as

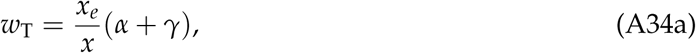

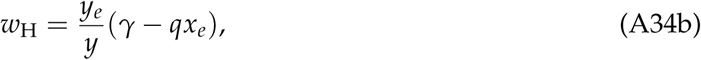

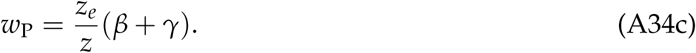

Replacing the ratios on the right side of these equations by the expressions in equation (A27) and using (from (A27a), (A33a), and (A33b))

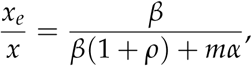

yields

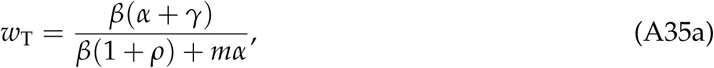

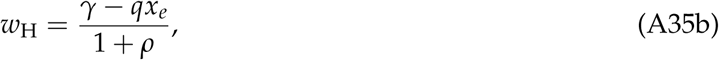

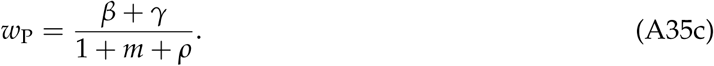

## Appendix B: Analysis of the Evolutionary Dynamics

The replicator dynamics (A31) has three trivial rest points at the corners of the simplex Δ: (*x, y, z*) = (1, 0, 0), (*x, y, z*) = (0, 1, 0), and (*x, y, z*) = (0, 0, 1). With slight abuse of notation, we denote these rest points by T, H, and P, respectively. In addition to analyzing the stability of the trivial rest points, our analysis consists in identifying the number, location, and stability of non-trivial rest points, and in how the phase portraits of our model depend on parameter values. Our analysis proceeds in six steps. First, we obtain convenient expressions for the pairwise comparison of the renormalized fitnesses in (A35) which provide the basis for much of the sub-sequent analysis (Section Pairwise fitness comparisons). Second, we show that the replicator dynamics (A31) has no interior rest point, that is, no rest point (*x, y, z*) in the interior of Δ, i.e., where *x* > 0, *y* > 0, and *z* > 0 (Section The replicator dynamics has no interior rest point). Thus, if the replicator dynamics has any rest points different from the trivial ones, these must be located on the edges of the simplex. Third, we investigate the dynamics along the three edges of the simplex Δ, thereby identifying how the number and location of the rest points on the edges of the simplex depend on the parameters of the model (Section Dynamics on the edges). This analysis provides us with much of the requisite information to determine the stability properties of all the rest points. Fourth, we complete the stability analysis for the non-trivial rest points identified in the third step (Section Stability analysis of the non-trivial rest points). Fifth, we summarize our results by identifying the five disjoint regions in our parameter space, each one characterized by qualitatively different phase portraits, shown in fig. 1 in the main text (Section Dynamical regions). Sixth, we provide for formal underpinnings for fig. 2 in the main text, showing how the five regions identified in the fifth step change as the parameters *q* and *ρ* change (Section Effects of varying *q* and *ρ* on the dynamical regions *ρ*). All together, these results provide us with a complete qualitative picture of the evolutionary dynamics of our model.

Throughout the following we write =_*s*_ to indicate that two expressions have the same sign (either +, −, or 0).

### Pairwise fitness comparisons

#### Comparison of *w*_P_ and *w*_T_

From (A35a) and (A35c) we obtain that *w*_P_ = *w*_T_ holds if and only if *β*(1 + *ρ*) = *mγ*:

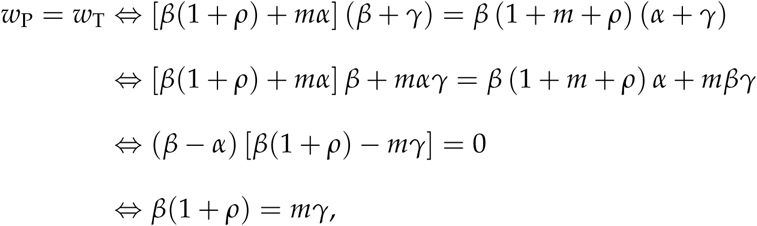

where the last equivalence follows from observing, first, that from (A33a) and (A33b) we have *β - α* = *λ*(1 − *e*) + *qy*_*e*_, and, second, that the latter expression is strictly positive as we have assumed *λ* > 0 and every steady-state satisfies *e* < 1 – unless we have *ρ* = 0 and *y* = 1, in which case the term *qy*_*e*_ = *qy* is strictly positive as we have assumed *q* > 0.

The same line of reasoning holds when we start with inequality rather than equality, showing

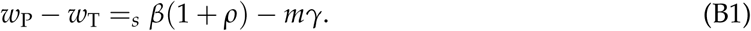

#### Comparison of *w*_P_ and *w*_H_

Using (A35b) and (A35c) we obtain

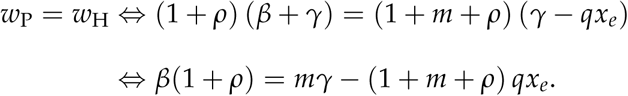

Similar reasoning implies:

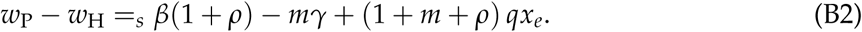

#### Comparison of *w*_T_ and *w*_H_

Using (A35a) and (A35b) we obtain

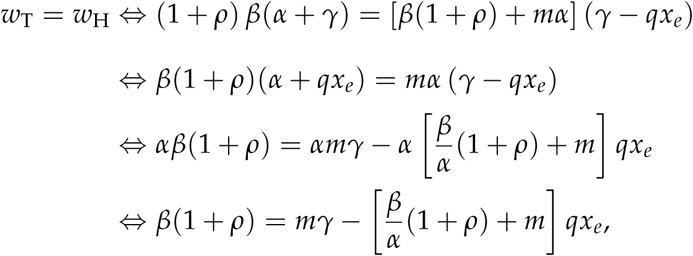

where we have used *α* > 0 (from (A33a), *e* ≥ *y*_*e*_, *q* < 1, and *e* > 0 for all (*x, y, z*) ∈ Δ) to obtain the last two equivalences. Similar reasoning implies that the sign of *w*_T_ − *w*_H_ coincides with the sign of 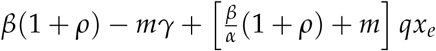 or:

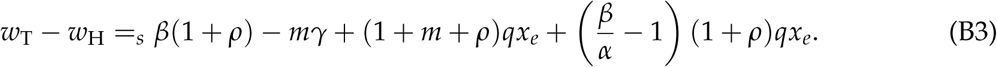

### The replicator dynamics has no interior rest point

If (*x, y, z*) is an interior rest point of the replicator dynamics, then the associated (*x*_*e*_, *y*_*e*_, *z*_*e*_) satisfies *x*_*e*_ > 0, *y*_*e*_ > 0, and *z*_*e*_ > 0, and we have *w*_P_ = *w*_T_ = *w*_H_. In particular, we must have *w*_P_ = *w*_T_ and *w*_P_ = *w*_H_. From (B1) and (B2) these equalities are equivalent to *β*(1 + *ρ*) = *mγ* and *β*(1 + *ρ*) = *mγ* − (1 + *m* + *ρ*) *qx*_*e*_. Substituting the first of these equalities into the second yields *qx*_*e*_ = 0. Because *q* > 0 holds, this contradicts *x*_*e*_ > 0. Therefore, no interior rest point exists. As a corollary, we also have that there are no closed orbits in the system (Strogatz, 1994, p. 180).

### Dynamics on the edges

#### TP-edge

On the TP-edge, the dynamics depend on how *m* compares to 1 + *ρ* and on how *λ* compares to the critical values

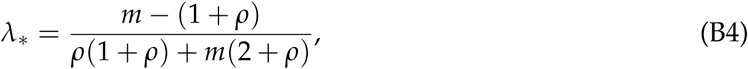

and

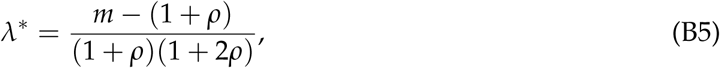

which for *m* > 1 + *ρ* satisfy *λ*_*_ < *λ*\*, in the following way (fig. B1):

**Figure B1:**
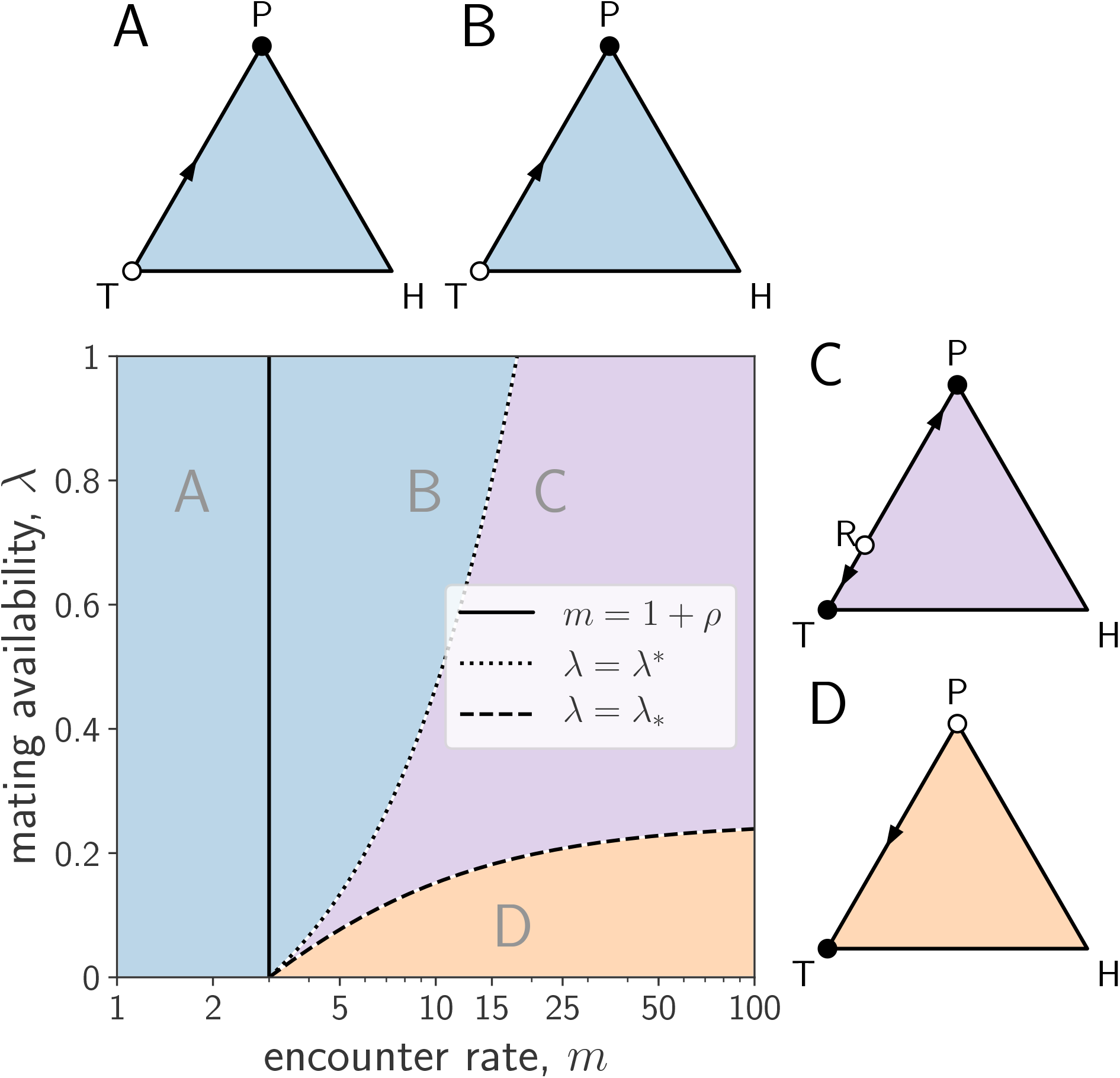
Evolutionary dynamics on the TP-edge. If *m* ≤ 1 + *ρ*, then P dominates T (A). The same is true if *m* > 1 + *ρ* and *λ* ≥ *λ*\* (B). If *λ*_*_ < *λ* < *λ*\*, there is bistability with both T and P being stable along the TP-edge (C). If *λ* ≤ *λ*_*_, T dominates P (D). Full (resp. empty) circles represent stable (resp. unstable) equilibria along the TP-edge. Parameters: *ρ* = 2, *q* = 0.5, *m* = 0.8 (A), 8 (B), 20 (C) or 15 (D), and *λ* = 0.75 (A, B, and C), or 0.05 (D).

1. If *m* ≤ 1 + *ρ*, then providing dominates trading, i.e., the dynamics on the TP-edge are unidirectional leading from T to P (fig. B1A).
2. If *m* > 1 + *ρ* and *λ*\* ≤ *λ*, then providing dominates trading (fig. B1B).
3. If *m* > 1 + *ρ* and *λ*_*_ < *λ* < *λ*\*, there is bistability, i.e., there exists a critical proportion of traders *x*_R_ ∈ (0, 1) such that R = (*x*_R_, 0, 1 − *x*_R_) is a rest point of the replicator dynamics and on the TP-edge the dynamics lead to P for *x* < *x*_R_ and to T for *x* > *x*_R_ (fig. B1C). For *λ* = 1 this critical proportion of traders is given by

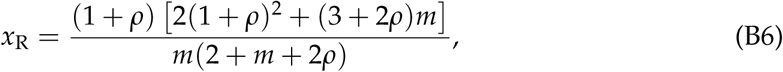

while for 0 < *λ* < 1 it is given by

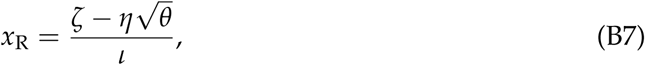

where

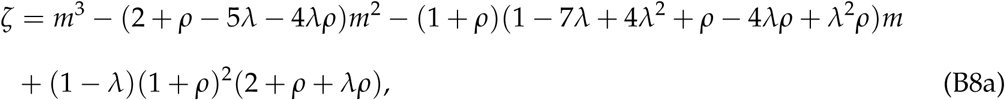

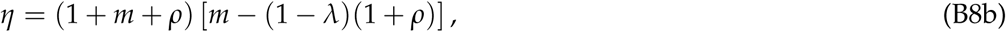

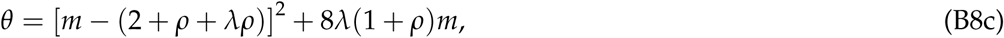

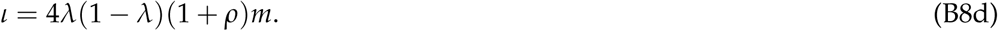
4. If *m* > 1 + *ρ* and *λ* ≤ *λ*_*_, then trading dominates providing, i.e., the dynamics on the TP-edge are unidirectional, leading from P to T (fig. B1D).

To prove the above claims, note that, as indicated in (B1), the sign of the payoff difference *w*_P_ − *w*_T_ coincides with the sign of *β*(1 + *ρ*) − *mγ*. On the TP-edge, *y* = 0 holds and hence *y*_*e*_ = 0 and *e* = *x*_*e*_ + *z*_*e*_ hold. Replacing the expressions for *β* and *γ* from their definitions (A33b) – (A33c) we thus obtain

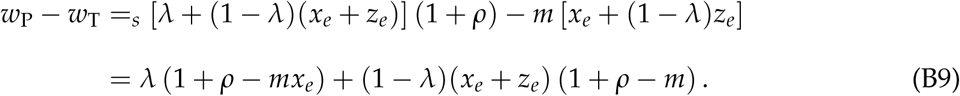

As both *x*_*e*_ and *z*_*e*_ are uniquely determined by *x* on the TP-edge, the latter explicitly as

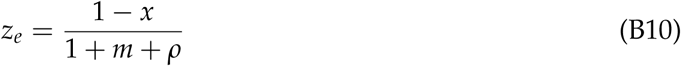

(by (A22c) and *z* = 1 − *x*) and the former by the unique solution to the equation (cf. (A21a))

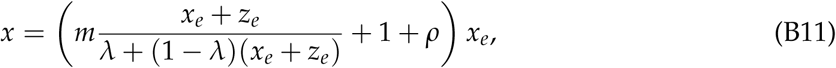

we may view the expression on the right side of (B9) as a function of *x* defined on the domain *x* ∈ [0, 1], that we denote by *h*(*x*).

For *m* ≤ 1 + *ρ*, the function *h*(*x*) is positive so that *w*_P_ − *w*_T_ > 0 holds for all *x* ∈ [0, 1]. This establishes the result for the first of the above cases.

In the remaining three cases we have *m* > 1 + *ρ*, which we may therefore impose as an assumption. We structure the argument for theses cases as follows: First, we show that *h*(*x*) is a decreasing function of *x*. Second, we assess how the extreme values *h*(0) and *h*(1) compare to zero. In particular, (i) if *h*(0) ≤ 0 then *h*(*x*) < 0 and hence *w*_P_ − *w*_T_ < 0 holds for *x* > 0 (trading dominates providing), (ii) if *h*(1) ≥ 0 then *h*(*x*) > 0 and hence *w*_P_ − *w*_T_ > 0 holds for 0 ≤ *x* < 1 (providing dominates trading), (iii) if *h*(1) < 0 < *h*(0) then there is bistability, as the fitness difference *w*_P_ − *w*_T_ is positive for *x* ∈ [0, *x*_R_) and negative for (*x*_R_, 1], where *x*_R_ is a root of *h*(*x*) such that *h*(*x*_R_) = 0.

To show that *h*(*x*) is decreasing in *x*, we consider the derivative of *h*(*x*), given by

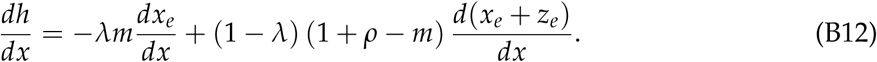

Using the inequality *m* > 1 + *ρ*, this is negative if both derivatives appearing on the right side of (B12) are positive. To show that this is the case, we differentiate both sides of the identity (B11) with respect to *x* to obtain

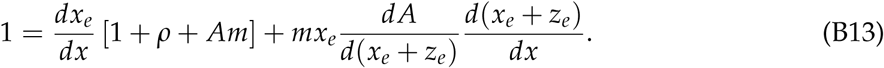

where we have used the abbreviation *A* = (*x*_*e*_ + *z*_*e*_)/(*λ* + (1 − *λ*)(*x*_*e*_ + *z*_*e*_)). Using *d*(*x*_*e*_ + *z*_*e*_)/*dx* = *dx*_*e*_/*dx* + *dz*_*e*_/*dx* and solving for *dx*_*e*_/*dx* we get

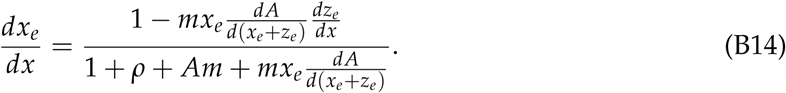

A straightforward calculation verifies that we have *dA*/*d*(*x*_*e*_ + *z*_*e*_) > 0. As we also have *dz*_*e*_/*dx* < 0 and *A* > 0, it follows from (B14) that *dx*_*e*_/*dx* > 0 holds. It remains to exclude the possibility that *d*(*x*_*e*_ + *z*_*e*_)/*dx* ≤ 0 in equation (B12). Towards this end, we observe that if *d*(*x*_*e*_ + *z*_*e*_)/*dx* ≤ 0 holds, then (B13) implies *dx*_*e*_/*dx* ≥ 1/(1 + *ρ* + *Am*). As *A* < 1 holds, we also have 1/(1 + *ρ* + *Am*) > 1/(1 + *ρ* + *m*), so that *dx*_*e*_/*dx* > 1/(1 + *ρ* + *m*). As *dz*_*e*_/*dx* = −1/(1 + *ρ* + *m*) it then follows that *d*(*x*_*e*_ + *z*_*e*_)/*dx* > 0 holds, yielding a contradiction. We conclude that *h*(*x*) is a decreasing function of *x*.

Next, we determine the sign of *h*(0). For *x* = 0 we have *x*_*e*_ = 0 and *e* = *z*_*e*_ = 1/(1 + *m* + *ρ*). Therefore,

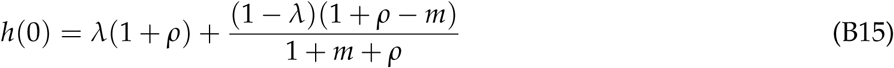

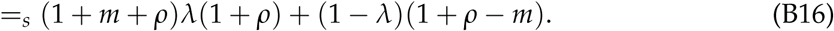

Consequently, the sign of *h*(0) coincides with the sign of *λ* − *λ*_*_, where *λ*_*_ is given by equation (B4).

In particular, the conditions *m* > 1 + *ρ* and *λ* ≤ *λ*_*_ ensure that *h*(*x*) is decreasing and that *h*(0) ≤ 0 holds. Consequently, under these conditions we have *h*(*x*) < 0 for *x* > 0, so that trading is dominant on the TP-edge. This establishes the fourth of the above claims.

It remains to consider *m* > 1 + *ρ* and *λ* > *λ*_*_. Here we have that *h*(*x*) is decreasing and *h*(0) > 0 holds. Therefore, if *h*(1) ≥ 0 holds, then providing is dominant on the TP-edge (i.e., we 40 are in the second of the above cases). Otherwise, i.e., if *h*(1) < 0 holds, then there exists a unique value 0 < *x*_R_ < 1 such that *h*(*x*_R_) = 0 holds and there is bistability on the TP-edge with the rest point corresponding to *x*_R_ separating the basins of attraction of T and P (i.e., we are in the third of the above cases). It remains to link the above conditions on the sign of *h*(1) to the conditions on *λ* appearing in our claims.

Consider the condition for *h*(1) ≥ 0, ensuring that providing is dominant along the TP-edge. As *x* = 1 implies *z*_*e*_ = 0, from (B9) we have

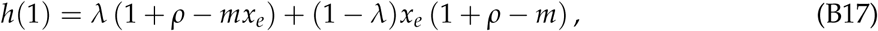

and from (B11) we have

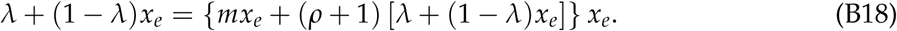

The unique positive solution to the quadratic implicitly defined by (B18) is

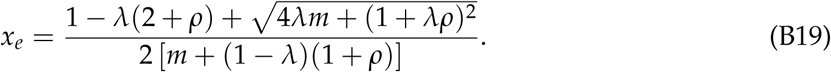

From equation (B17), and noting that *m* > (1 + *ρ*)(1 − *λ*) holds (since we assumed that *m* > 1 + *ρ* holds), the condition *h*(1) ≥ 0 can then be written as

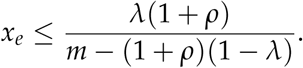

Substituting equation (B19) into the above expression, rearranging, and simplifying, we obtain that *h*(1) ≥ 0 is equivalent to

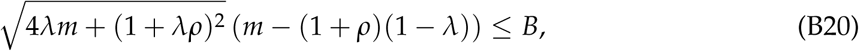

where we have defined

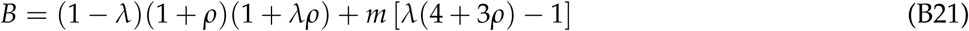

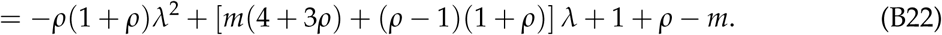

The expression on the left hand side of (B20) is positive. *B* can be either negative or non-negative, depending on parameter values. If *B* is negative, condition (B20) cannot hold, and hence *h*(1) < 0 must hold. If *B* is non-negative, taking squares of both sides of (B20) and simplifying shows that *λ* ≥ *λ*\* (where *λ*\* is given by equation (B5)) is a necessary and sufficient condition for (B20) (and hence *h*(1) ≥ 0) to hold. In particular, no matter the sign of *B, h*(1) < 0 holds if *λ* < *λ*\*.

To show that *h*(1) ≥ 0 holds if *λ* > *λ*\*, it remains to exclude the possibility that *B* is negative when *λ* > *λ*\*. From (B21), a necessary condition for *B* to be negative is that

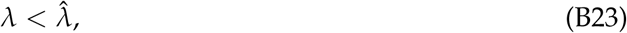

where

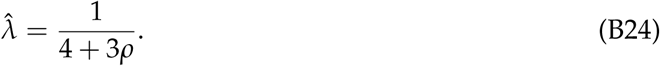

We could have the following two scenarios:

First, 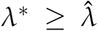. In this case, *λ* ≥ *λ*\* implies that condition (B23) is violated, so that *B* is non-negative.

Second, 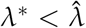. Then, if 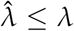 also holds, condition (B23) is violated and *B* is non-negative. It remains to show that *B* is non-negative if 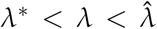 holds. To do so, note that *B* can be written as a quadratic function in *λ* (equation (B22)), *B*(*λ*). In this case, *B*(*λ*) has two roots in the positive axis, and *B*(0) < 0 and lim_*λ*→∞_ *B*(*λ*) < 0 hold for *m* > 1 + *ρ*. Since *B*(*λ*\*) > 0 and 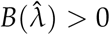 hold for *m* > 1 + *ρ*, this implies that *B*(*λ*) is positive in the whole interval 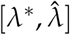.

We conclude that *h*(1) ≥ 0 holds if *λ* ≥ *λ*\* and that *h*(1) < 0 holds if *λ*_*_ < *λ* < *λ*\*.

To find the value *x*_R_ ∈ (0, 1) such that *h*(*x*_R_) = 0 holds when there is bistability, first assume that *λ* = 1 holds. Then the right hand side of (B9) reduces to 1 + *ρ* − *mx*_*e*_, so that *h*(*x*_R_) = 0 is equivalent to

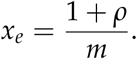

Replacing equation (A22a) into this expression (with *λ* = 1, *x* = *x*_R_, *y* = 0, *z* = 1 − *x*_R_), solving for *x*_R_, and simplifying, yields the expression for *x*_R_ given in equation (B6).

Assuming now that 0 < *λ* < 1, *h*(*x*_R_) = 0 is equivalent to

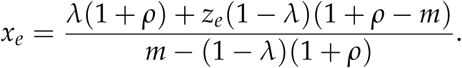

Replacing equation (A22a) and equation (B10) into this expression (with *x* = *x*_R_, *y* = 0, *z* = 1 − *x*_R_), solving for *x*_R_, and simplifying, yields the expression for *x*_R_ given in equation (B7).

#### HP-edge

On the HP-edge, the dynamics depend on how *λ* compares to the critical value *λ*_*_ given in equation (B4) in the following way (fig. B2):

1. If *λ* ≥ *λ*_*_, then providing dominates withholding, i.e., the dynamics on the HP-edge are unidirectional and lead from H to P (fig. B2A).
2. If *λ* < *λ*_*_, then providers can invade H, withholders can invade P, and there exists one further rest point S = (0, 1 − *z*_S_, *z*_S_) on the HP-edge (fig. B2B). The proportion of providers at this rest point is given by

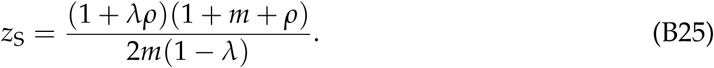

**Figure B2:**
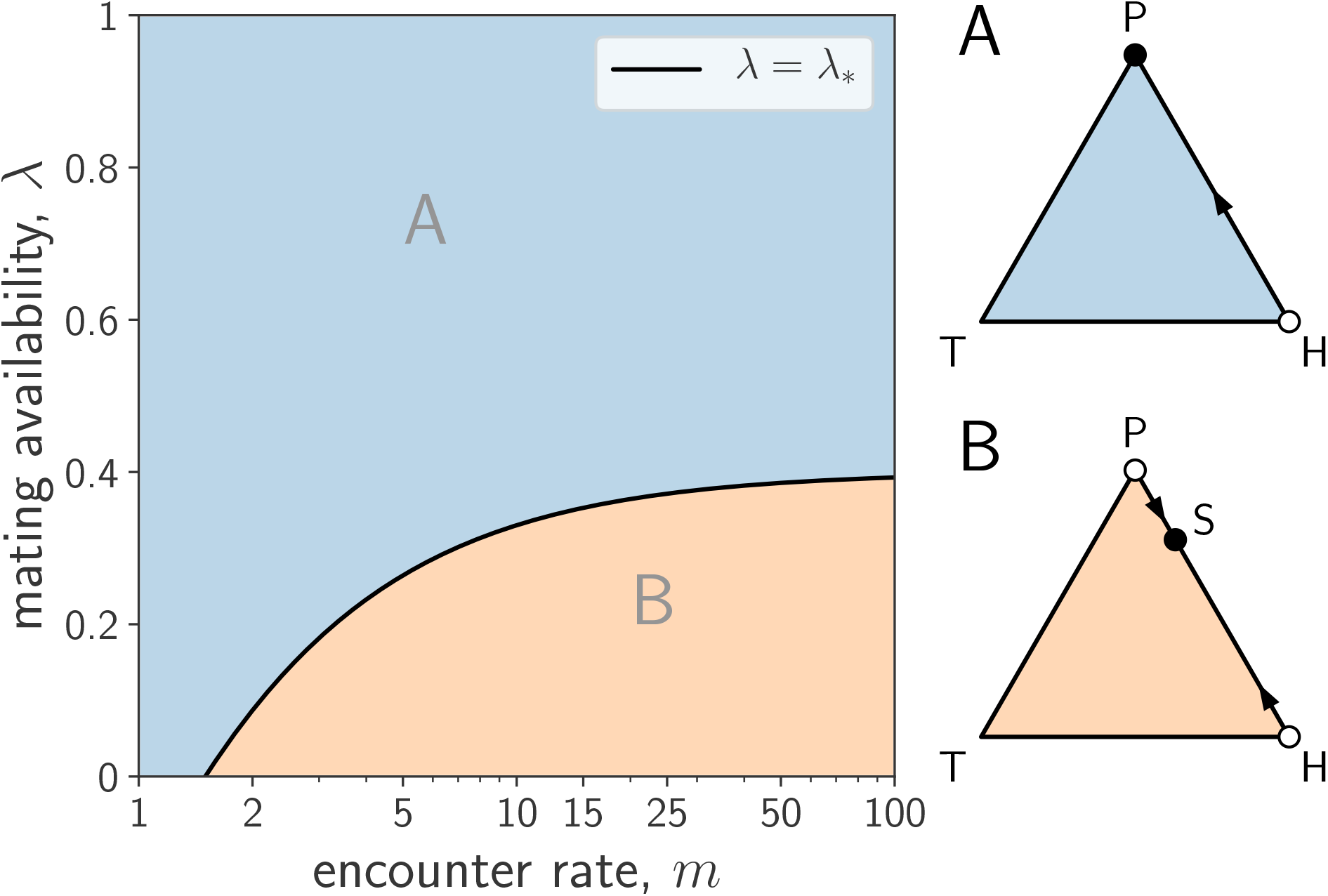
Evolutionary dynamics on the HP-edge. If *λ* ≥ *λ*_*_, providing dominates withholding (A). If *λ* < *λ*_*_, traders invade P, providers invade T, and the two strategies coexist at a polymorphic equilibrium S (B). Full (resp. empty) circles represent stable (resp. unstable) equilibria along the HP-edge. Parameters: *ρ* = 0.5, *q* = 0.5, *m* = 5 (A), or 20 (B), and *λ* = 0.7 (A), or 0.2 (B).

To show this, note that on the HP-edge, *x* = 0 and hence *x*_*e*_ = 0 holds. Therefore, as indicated in (B2), the sign of the payoff difference *w*_P_ − *w*_H_ coincides with the sign of *β*(1 + *ρ*) − *mγ*. Replacing the expressions for *β* and *γ* from their definitions (A33b) – (A33c) and using *e* = *y*_*e*_ + *z*_*e*_ we thus obtain

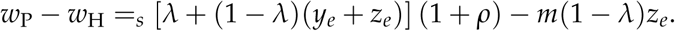

Replacing the expressions for *y*_*e*_ and *z*_*e*_ (equation (A22b) and (A22c)) with *y* = 1 − *z*, and simplifying, we obtain

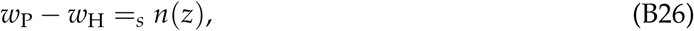

where

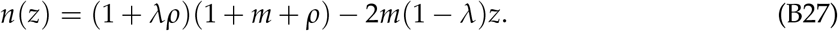

Since *n*(0) = (1 + *λρ*)(1 + *m* + *ρ*) is positive, the linear function *n*(*z*) (and hence the payoff difference (B26)) is either positive for all *z* ∈ [0, 1), or has a single sign change at some *z*_S_ ∈ (0, 1) on the HP-edge.

A necessary and sufficient condition for *n*(*z*) to change sign is that *n*(1) < 0 holds. This condition is satisfied if and only if *λ* < *λ*_*_, where *λ*_*_ is given by equation (B4). In this case, the point *z*_S_ at which the direction of selection changes is found by solving the linear equation *n*(*z*_S_) = 0 for *z*_S_. Otherwise, if *λ* ≥ *λ*_*_, *n*(0) ≥ 0 holds, and the sign of *n*(*z*) (and hence of the payoff difference (B26)) is positive in the relevant interval. This establishes our claims.

#### TH-edge

On the TH-edge, the dynamics depend on how *λ* compares to the critical value

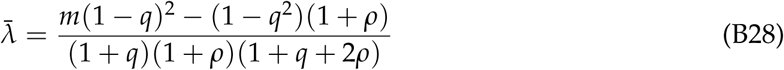

in the following way (fig. B3):

1. If 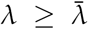, then trading dominates withholding, i.e., the dynamics on the TH-edge are unidirectional and lead from H to T (fig. B3A).
2. If 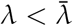, then traders can invade H, withholders can invade T, and there exists one further rest point Q = (*x*_Q_, 1 − *x*_Q_, 0) on the TH-edge (fig. B3B). The proportion of traders *x*_Q_ at this rest point is given by

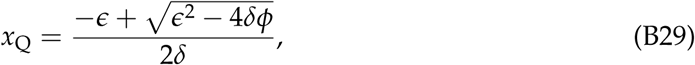

**Figure B3:**
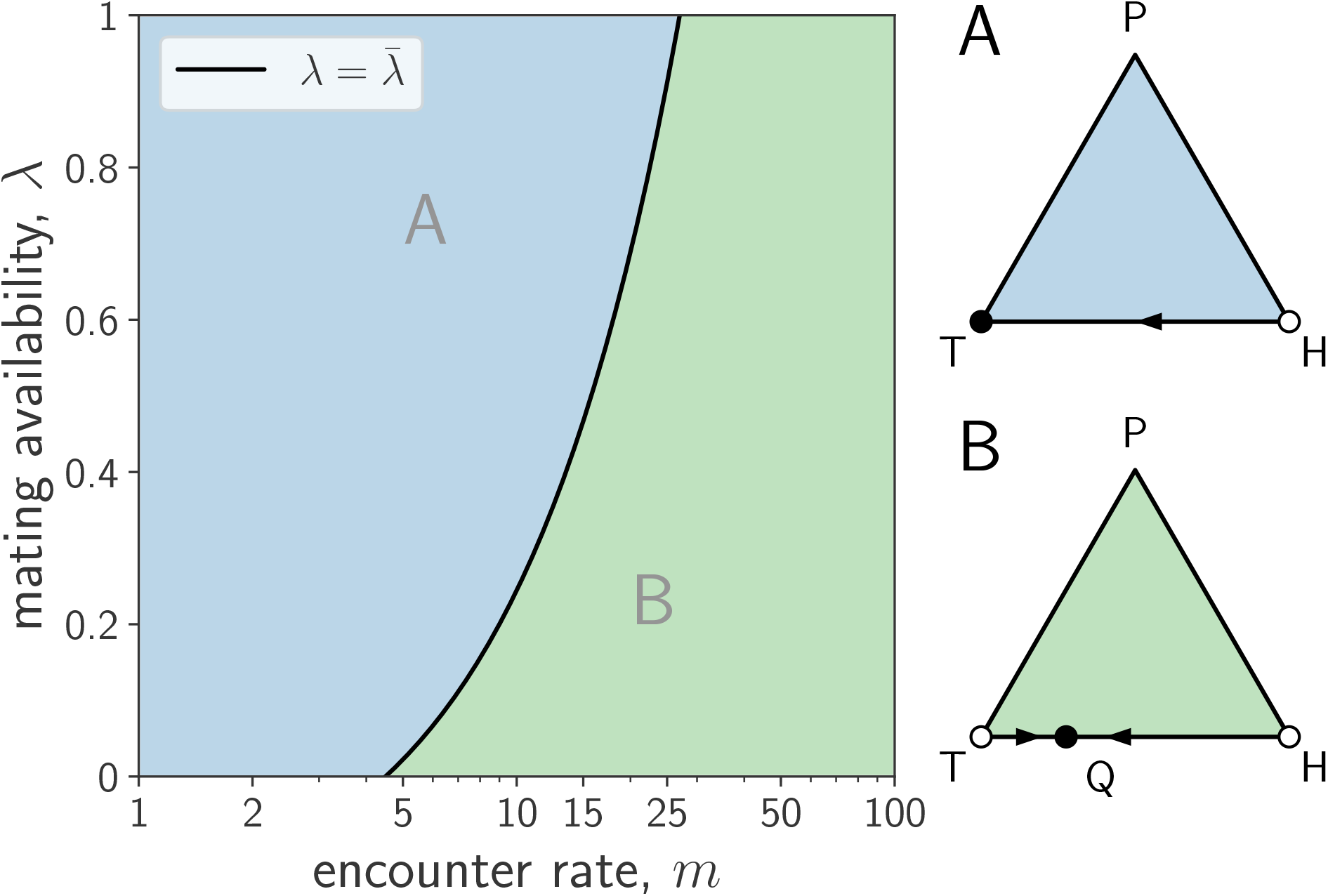
Evolutionary dynamics on the TH-edge. If 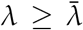, trading dominates withholding (A) If 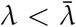, rare traders invade H, rare withholders invade T, and the two strategies coexist at a polymorphic equilibrium Q (B). Parameters: *ρ* = 0.5, *q* = 0.5, *m* = 5 (A), or 20 (B), and *λ* = 0.7 (A), or 0.2 (B).

where

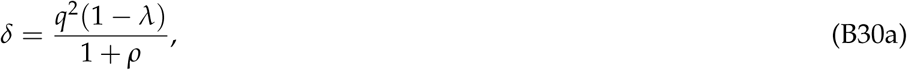

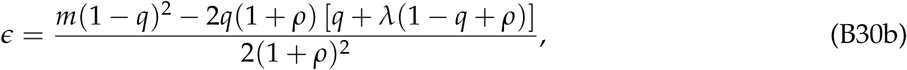

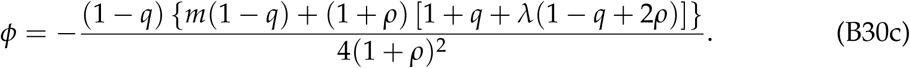

Moreover, *x*_Q_ is decreasing in *m* and tends to 1/2 as *m* grows large.

To show this, note that on the TH-edge, *z* = 0 and hence *z*_*e*_ = 0 holds. Setting *z*_*e*_ = 0 in equation (A32) we obtain

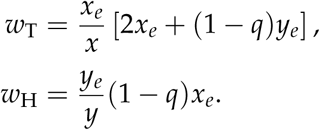

Replacing the expression for *y*_*e*_ (equation (A22b)) with *y* = 1 − *x* into the above payoffs and simplifying, we obtain

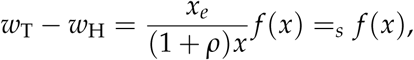

as *x*_*e*_/[(1 + *ρ*)*x*] is always positive for *x* ∈ (0, 1), and where we have defined

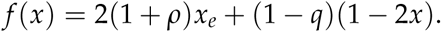

Along the TH-edge, *x*_*e*_ is given by equation (A22a), with

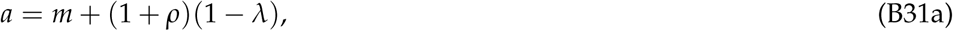

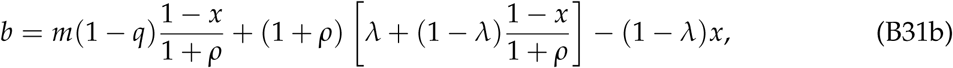

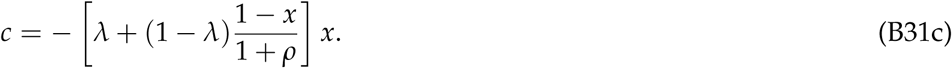

It is clear that *f* (0) = 1 − *q* > 0, and that the roots of *f* (*x*) satisfy

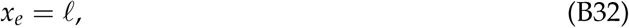

where we have used the abbreviation

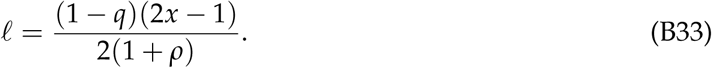

In particular, since *x*_*e*_ ≥ 0 and *ℓ* < 0 always holds if *x* < 1/2, it must be that roots of *f* (*x*) can only exist in the interval [1/2, 1].

Substituting (A22a) into (B32) we obtain

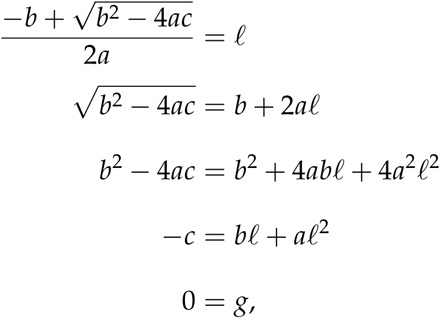

where we defined

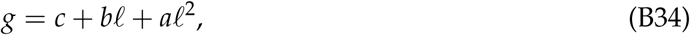

which can be viewed as a function of *x, g*(*x*). Note that the roots of *f* (*x*) and *g*(*x*) coincide. Moreover, since *b* and *ℓ* are linear in *x* and *c* is quadratic in *x, g*(*x*) is a quadratic function of *x* that can be rewritten as

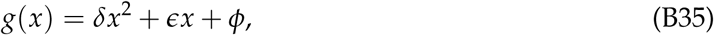

for real coefficients *δ, ϵ*, and *ϕ*. Replacing the expressions for *a, b, c* (given in (B31)) and the expression for *ℓ* (given in equation (B33)), into (B34) and simplifying, we obtain the values of these coefficients as given by (B30). Since *δ* > 0 and *ϕ* < 0 always hold, and by Descartes’ rule of signs, *g*(*x*) (and hence *f* (*x*)) has exactly one positive root *x*_Q_, given by equation (B29). Since *g*(0) = *ϕ* < 0, a necessary and sufficient condition for *x*_Q_ < 1 is that *g*(1) > 0 holds. Substituting *x* = 1 into equation (B35) and simplifying, we get

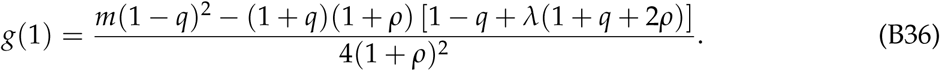

From this expression, it is immediate that a necessary and sufficient condition for *g*(1) > 0 (and hence *f* (1) < 0) is that the numerator of (B36) is positive, which obtains if and only if 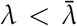, where 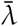 is given by (B28). In this case, and since *f* (0) > 0, *f* (*x*) is positive for *x* ∈ [0, *x*_Q_) and negative for (*x*_Q_, 1]. This establishes that the condition 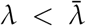 ensures that traders and withholders invade each other and coexist at an equilibrium frequency *x*_Q_ given by equation (B29). Otherwise, if 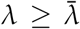, then *g*(1) ≤ 0 and there is no root of *g*(*x*) or *f* (*x*) in the interval (0, 1). In this case, it follows that *f* (*x*) is positive for all *x* ∈ [0, 1]. This establishes that trading dominates withholding for 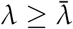.

It remains to show that the proportion of traders *x*_Q_ at the equilibrium Q is decreasing in the mate encounter rate *m* and tends to 1/2 as *m* grows large. To do so, first note that, from equation (B35), *x*_Q_ is given implicitly by

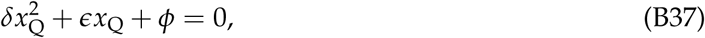

where *δ, ϵ*, and *ϕ* are as given in equation (B30). Differentiating implicitly with respect to *m* and simplifying we obtain

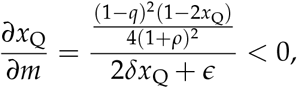

where inequality follows from the fact that the denominator is positive, and that *x*_Q_ > 1/2 holds (as shown after equation (B33)). This establishes the monotonic decrease of *x*_Q_ with respect to *m*.

To obtain the limit result, divide both sides of equation (B37) by *ϵ*, take the limit of both sides when *m* → ∞, and simplify to obtain lim_*m*→∞_ *x*_Q_ = 1/2.

### Stability analysis of the non-trivial rest points

The previous analysis has identified three non-trivial rest points located on the edges of the simplex: Q (on the TH-edge), R (on the TP-edge), and S (on the HP-edge). Here, we discuss the local stability of these rest points.

#### Q is a sink

Suppose that the rest point Q, located on the TH-edge, exists. From the analysis in Section TH-edge, this rest point is stable along the TH-edge as it is attracting from both T and H.

Moreover, Q is also attracting for neighboring points in the interior of the simplex. To show this, we begin by noting that at Q the fitnesses of traders and withholders are equal, i.e., *w*_T_ = *w*_H_ holds. By (B3) this implies

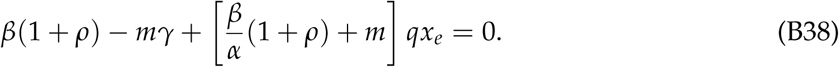

Since *α* and *β*, defined in (A33), are positive and at Q we also have *x* > 0 and hence *x*_*e*_ > 0, (B38) implies *β*(1 + *ρ*) − *mγ* < 0. By (B1) this is the condition for *w*_P_ < *w*_T_ to hold. We then have that at Q the fitnesses of the three strategies satisfy *w*_T_ = *w*_H_ > *w*_P_, establishing our claim.

We conclude that Q is a sink. In particular, it is stable.

#### R is saddle

Suppose that the rest point R, located on the TP-edge, exists. From the analysis in Section TP-edge, this rest point is unstable along the TP-edge as it is repelling from both T and P.

Moreover, R is attracting for neighboring points in the interior of the simplex. To show this, we begin by noting that at R the fitnesses of traders and providers are equal, i.e., *w*_T_ = *w*_P_ holds. By (B1) this implies *β*(1 + *ρ*) − *mγ* = 0. Since *q* > 0 and at R we have *x* > 0 and hence *x*_*e*_ > 0, this implies *β*(1 + *ρ*) − *mγ* + (1 + *m* + *ρ*)*qx*_*e*_ > 0. By (B2) this is the condition for *w*_P_ > *w*_H_ to hold. We then have that at R the fitnesses of the three strategies satisfy *w*_T_ = *w*_P_ > *w*_H_, establishing our claim.

Hence, R is a saddle. In particular, it is unstable.

#### S is a saddle

Suppose that the rest point S, located on the HP-edge, exists. From the analysis in Section HP-edge, this rest point is stable along the HP-edge as it is attracting from both H and P.

At S we have *w*_H_ = *w*_P_ and, further, *x*_*e*_ = 0 because *x* = 0 holds. By equations (B1) and (B2) this implies *w*_H_ = *w*_P_ = *w*_T_. Hence, we cannot use an argument similar to the one given in Sections Q is a sink and R is saddle to infer whether or not S is attracting for neighboring points in the interior of the simplex. We therefore resort to center manifold theory (Kuznetsov, 2013) to show that S is a saddle point. Throughout the following argument, we will make use of the fact that the rest point S only exists if *λ* < *λ*_*_ holds (Section HP-edge) and that *λ*_*_ < 1 holds (cf. (B4)), so that we may assume *λ* < 1.

As a first step, we observe that by using the identity *z* = 1 − *x* − *y* the fitnesses *w*_T_, *w*_H_, and *w*_P_ as given in equations (A35) can be expressed as functions of *x* and *y*. The evolutionary dynamics (A31) can then be reduced to the two-dimensional system

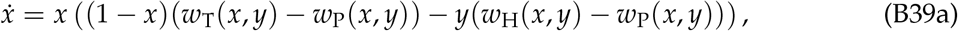

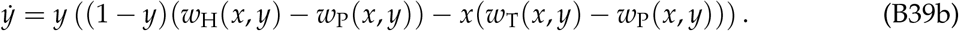

In terms of this system our interest is in determining the stability of the rest point (0, *y*\*), where *y*\* = 1 − *z*_S_ and *z*_S_ is given by equation (B25). The Jacobian of the dynamic at this rest point:

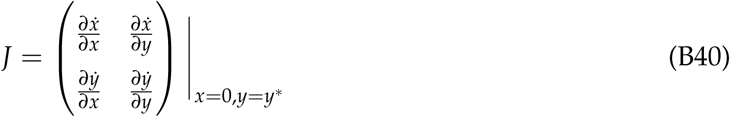

takes the form

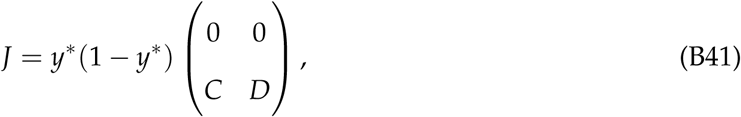

where

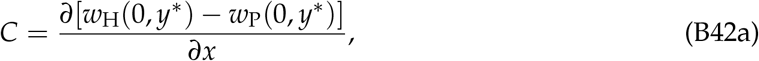

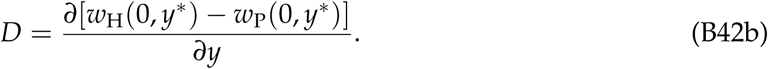

To obtain this result from (B39), we have used the fact that *w*_H_(0, *y*\*) = *w*_P_(0, *y*\*) = *w*_T_(0, *y*\*) holds.

The argument demonstrating the stability of the rest point S along the HP-edge in Section HP-edge implies that *w*_H_(0, *y*) − *w*_P_(0, *y*) is linear and decreasing in *y*. (In terms of the function *n* defined in equation (B27), we have *w*_H_(0, *y*) − *w*_P_(0, *y*) = −*n*(1 − *y*)/(1 + *ρ*)((1 + *m* + *ρ*)), with the inequality *λ* < 1 implying that *n*(1 − *y*) is increasing *y*.) Thus, we have *D* < 0. Consequently, the two eigenvalues of *J* are given by *µ*_1_ = *y*\*(1 − *y*\*)*D* < 0 and *µ*_2_ = 0 with associated eigenspaces *E*_1_ and *E*_2_ given by the scalar multiples of the eigenvectors *e*_1_ = (0, 1) and *e*_2_ = (1, −*C*/*D*). Note that the eigenspace *E*_1_ corresponds to movements along the HP-edge, so that the negativity of the eigenvalue *µ*_1_ reflects the stability of the dynamic along that edge. Center manifold theory asserts that there exists an invariant manifold of the dynamic that is tangent to the eigenspace *E*_2_ associated with the eigenvalue *µ*_2_ = 0 at the rest point (0, *y*\*). Further, the stability properties of the rest point are determined by the stability properties of the dynamic along this so-called center manifold. In our case only displacements from the rest point into the interior of the simplex are relevant. We now show that for a sufficiently small displacement onto the center manifold the trajectory starting from such an initial condition will lead away from the HP-edge, indicating that S is a saddle.

Continuing to use the identity *z* = 1 − *x* − *y* we can view the expressions appearing on the right sides of equations (B1) – (B3) as functions of *x* and *y*:

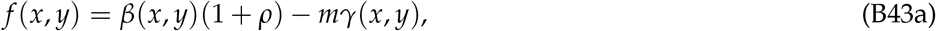

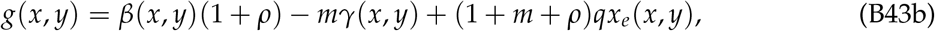

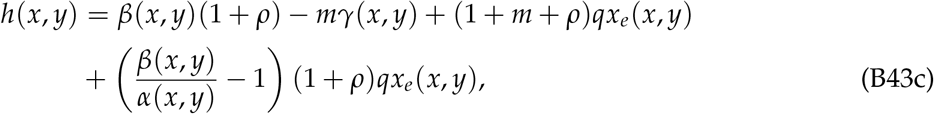

From equations (B1) – (B3) and *w*_H_(0, *y*\*) = *w*_P_(0, *y*\*) = *w*_T_(0, *y*\*) we have *f* (0, *y*\*) = *g*(0, *y*\*) = *h*(0, *y*\*) = 0.

The functions *f* (*x, y*), *g*(*x, y*), and *h*(*x, y*) are well defined and continuously differentiable on a neighborhood of the rest point (0, *y*\*). Further, appealing to the same arguments as the one leading up to equation (B27) in Section HP-edge we have that the functions defined in (B43) satisfy

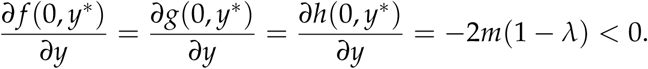

Therefore, the implicit function theorem yields the existence of continuously differentiable functions *y* ^*f*^ (*x*), *y*^*g*^ (*x*), and *y*^*h*^ (*x*), uniquely defined on some interval [0, *ϵ*), satisfying

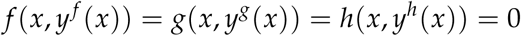

on that interval as well as *y* ^*f*^ (0) = *y*^*g*^ (0) = *y*^*h*^ (0) = *y*\*. Further, the derivatives of these functions at *x* = 0 are given by

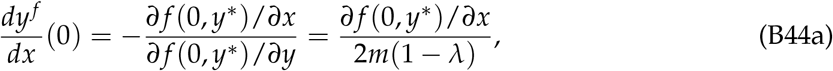

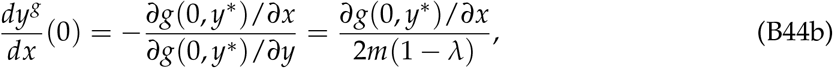

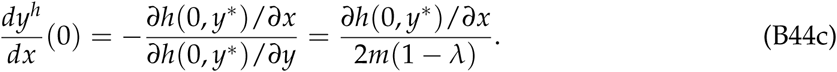

As *g*(*x, y*) differs from *w*_H_(*x, y*) − *w*_P_(*x, y*) only by a non-zero multiplicative constant, we also have

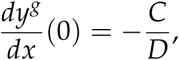

indicating that the center manifold is tangent to the graph of the function *y*^*g*^ at the rest point (0, *y*\*). Provided that

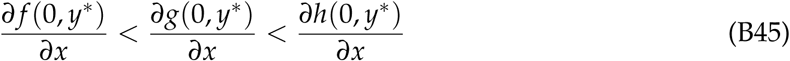

holds, it follows that for sufficiently small *x*^*c*^ > 0 a point (*x*^*c*^, *y*^*c*^) on the center manifold satisfies *y* ^*f*^ (*x*^*c*^) < *y*^*c*^ < *y*^*h*^ (*x*^*c*^) and therefore *f* (*x*^*c*^, *y*^*c*^) < 0 < *h*(*x*^*c*^, *y*^*c*^). From (B1) and (B3) it then follows that we have *w*_T_(*x*^*c*^, *y*^*c*^) > *w*_P_(*x*^*c*^, *y*^*c*^) and *w*_T_(*x*^*c*^, *y*^*c*^) > *w*_H_(*x*^*c*^, *y*^*c*^), implying that the population share *x* is increasing in a trajectory starting from (*x*^*c*^, *y*^*c*^).

To complete the argument, it remains to establish the inequalities in (B45). From (B43) we have *g*(*x, y*) − *f* (*x, y*) = (1 + *m* + *ρ*)*qx*_*e*_(*x, y*). Therefore, as (1 + *m* + *ρ*)*q* > 0 holds, the first inequality in (B45) holds if ∂*x*_*e*_(0, *y*\*)/∂*x* > 0. To see that this is true, we find it convenient to use implicit differentiation on (A26) to obtain

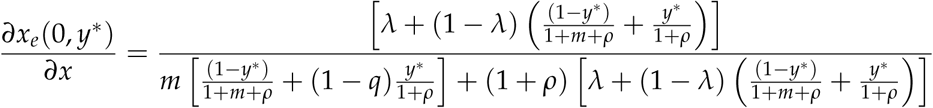

and observe that both numerator and denominator of the expression on the right side are positive. Similarly, we have

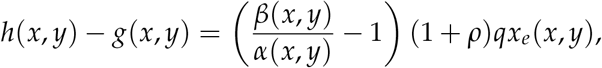

(*β*(*x, y*))

so that the second inequality in (B45) holds if the partial derivative of this expression with respect to *x* evaluated at (0, *y*\*) is positive. Applying the product rule, the derivative in question is given by

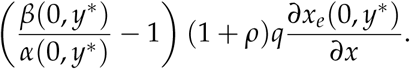

As *β*(*x, y*) > *α*(*x, y*) > 0 holds and we have already established ∂*x*_*e*_(0, *y*\*)/∂*x* > 0, this delivers the desired result.

### Dynamical regions

Here we build on the characterization of the dynamics on the edges from Section Dynamics on the edges to first establish in Section Co-existence of non-trivial rest points that, for any given values of the parameters 0 < *q* < 1 and *ρ* ≥ 0, and for generic values of the parameters 0 < *λ* ≤ 1 and *m* > 0, and five different scenarios for the co-existence of the rest points R, S, and Q arise. These are (i) none of these rest points exists, (ii) only ther est point R exists, (iii) only the rest point S exists, (iv) the rest points R and Q co-exist, and (v) the rest points S and Q co-exist. For each of these scenarios, the stability properties of the other three rest points (T, H, and P) are immediate from Section Dynamics on the edges, while the stability of whichever of the rest points R, S, and Q that exists has been established in Section Stability analysis of the non-trivial rest points. Combining this with the observation that there are no interior rest points or closed orbits (Section The replicator dynamics has no interior rest point) this provides us with a complete picture of the qualitative properties of the dynamics in each of the five different scenarios that we present in Section Characterization of the dynamics. Finally, Section Characterization of the dynamical regions in the main text characterizes the five different dynamical scenarios in terms of the inequality relationships that we employ in the main text.

#### Co-existence of non-trivial rest points

The existence of the non-trivial rest points depends on how *λ* compares to the critical values *λ*_*_, *λ*\*, and 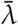 (respectively given by equations (B4), (B5), and (B28)). For given values of the parameters 0 < *q* < 1 and *ρ* ≥ 0 we consider these critical values as functions of *m* (fig. B4) and write

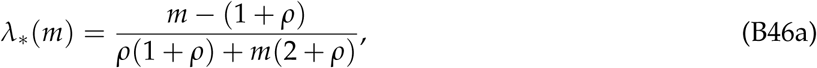

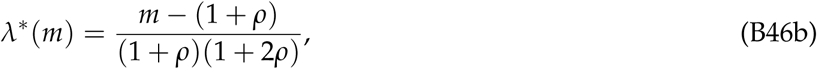

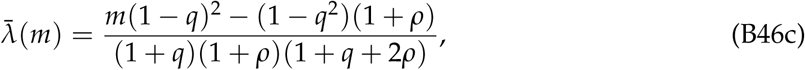

All these three functions are increasing in *m*. Moreover, *λ*_*_ is asymptotic to

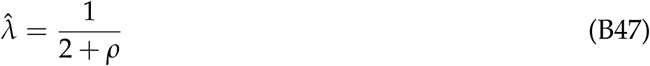

as *m* grows large; *λ*_*_ and *λ*\* are equal to zero at a critical value of *m* given by

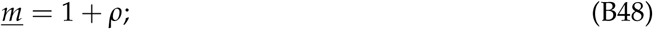

and 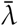 is equal to zero at a critical value of *m* given by

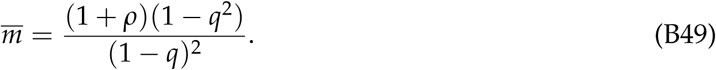

Since (1 − *q*^2^)/(1 − *q*)^2^ > 1 holds for 0 < *q* < 1, these critical values of *m* satisfy 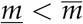.

It was already noted in Section TP-edge that for *m* > *<u>m</u>*, the inequality *λ*_*_(*m*) < *λ*\*(*m*) holds. As 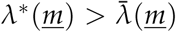 holds and it is easily verified that 0 < *q* < 1 implies that the derivatives of *λ*\* and 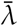 with respect to *m* satisfy 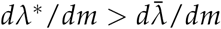, we also have the inequality 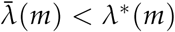 for all *m* ≥ *<u>m</u>*. It remains to investigate how 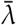 and *λ*_*_ compare.

**Figure B4:**
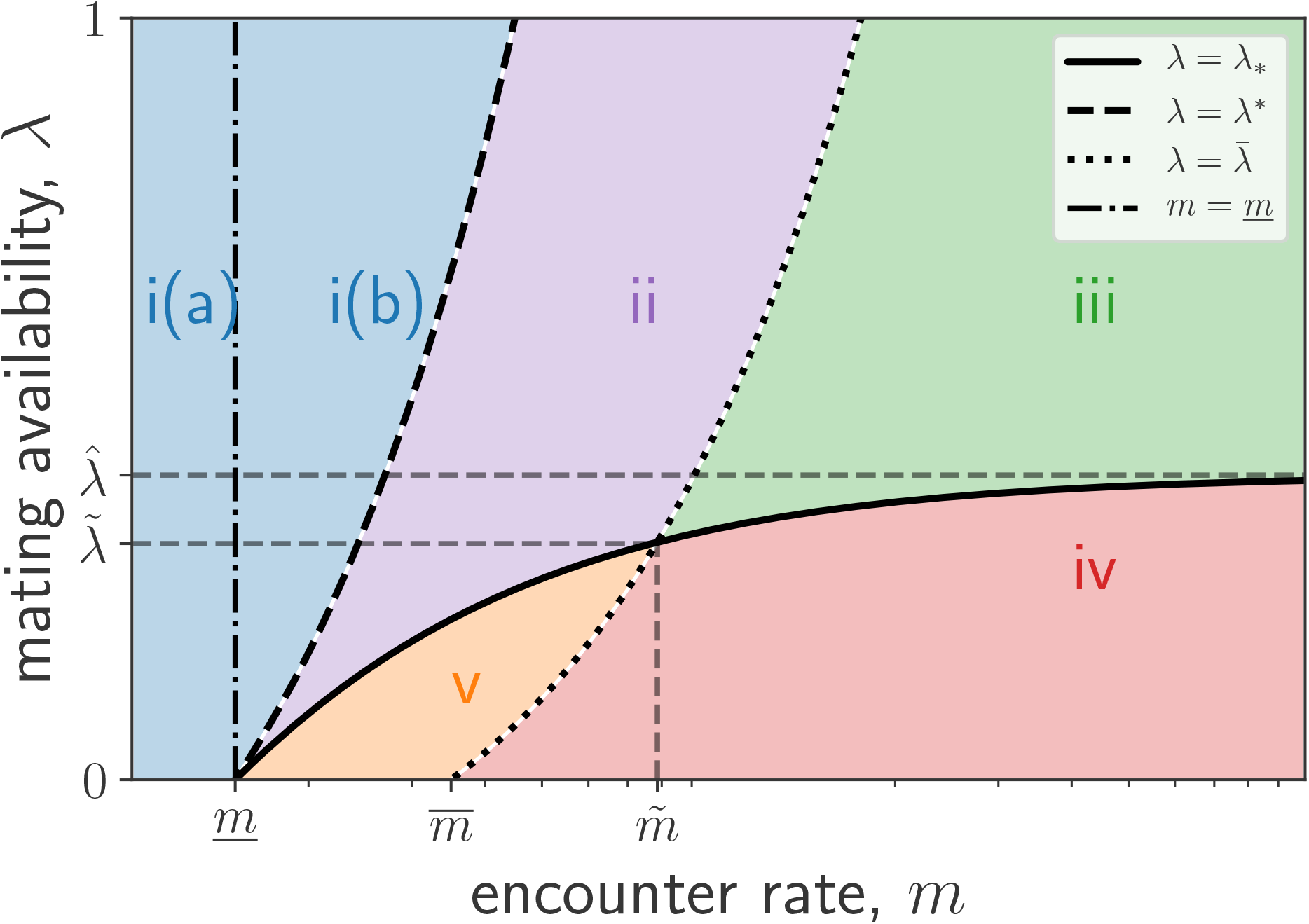
The five disjoint and non-empty regions into which the parameter space can be partitioned. The precise shape of these regions depends on the values of the parameters *ρ* and *q*, but the general picture is qualitatively the same. Parameters: *ρ* = 0.5, *q* = 0.4.

Consider the difference 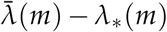 for 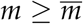. First, note that 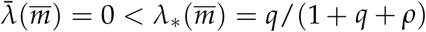, and hence 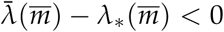. Second, we have 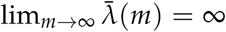, and lim_*m*→∞_ *λ*_*_(*m*) = 1/(2 + *ρ*), so that the difference 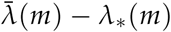 is positive when *m* is large. We then have that 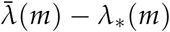 has an odd number of sign changes in 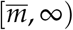. From (B46a) and (B46c), we have that 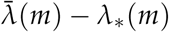 also satisfies

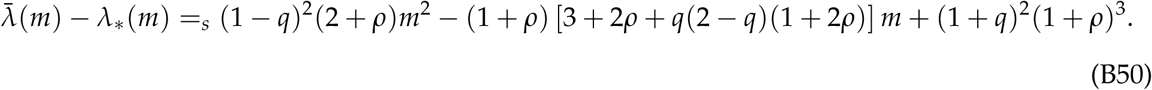

Denote the quadratic in *m* on the right hand side of the above expression by *p*(*m*). By Descartes’ rule of signs, *p*(*m*) and hence 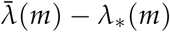 has either zero or two sign changes in the interval [0, ∞). Since we have established that 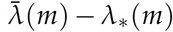 has an odd number of sign changes in 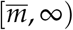, it must be that 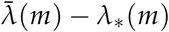 has two positive roots: one in the interval 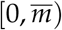 and another in the interval 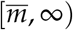. Moreover, at this latter root, 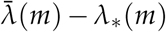 changes sign from negative to positive. Consequently, there exists a uniquely determined value 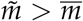 such that for 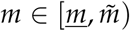 we have 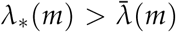, for 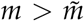 we have 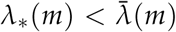, and for 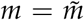 we have 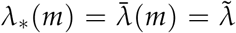, where 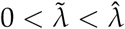 (fig. B4).

The properties of the functions *λ*_*_(*m*), *λ*\*(*m*), and 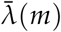 established above imply that the set of feasible values for the parameters *m* > 0 and 0 < *λ* ≤ 1 can be partitioned into five disjoint and non-empty regions as follows (where we ignore the non-generic cases in which one of the inequalities involving *λ* holds as an equality; fig. B4):

i. (a) *m* ≤ *<u>m</u>* or (b) *m* > *<u>m</u>* and *λ*\*(*m*) < *λ*.
ii. *m* > *<u>m</u>* and 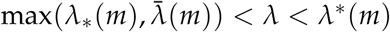.
iii. *m* > *<u>m</u>* and 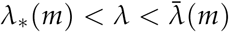.
iv. *m* > *<u>m</u>* and 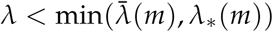.
v. *m* > *<u>m</u>* and 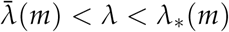.

In the first of these regions none of the non-trivial rest points R, S, and Q exists. To see this, consider case (a) first. Here *λ*_*_(*m*) and 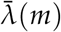 are both non-positive, so that the inequalities *λ* ≥ *λ*_*_(*m*) and 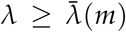 are implied by *λ* > 0. The results from Section Dynamics on the edges then imply that none of the non-trivial rest points R, S, and Q exists. In case (b) we have *λ*\*(*m*) > *λ*_*_(*m*) and 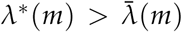 is not only strictly greater than *λ*\*(*m*), but also strictly greater than *λ*_*_(*m*) and 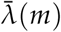. The results from Section Dynamics on the edges then imply that in this region, too, none of the non-trivial rest points R, S, and Q exists.

In the second region, the inequality 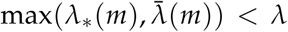 implies that neither of the rest points S and Q exist, whereas the inequalities *λ*_*_(*m*) < *λ* < *λ*\*(*m*) imply that the rest point R exists. Thus, in this region R is the only non-trivial rest point.

In the third region, we again have *λ*_*_(*m*) < *λ* < *λ*\*(*m*), so that the rest point R exists, whereas the rest point S does not exist. The additional inequality 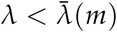 implies that, in addition to R, the rest point Q exists.

In the fourth region, the inequality *λ* < *λ*_*_(*m*) implies (as *λ*_*_(*m*) < *λ*\*(*m*) holds) that the rest point R does not exist, whereas the rest point S exists. From the inequality 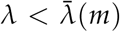, the rest point Q exists, too, so that in this region the rest points Q and S co-exist.

In the fifth region, the inequality again implies that the rest point R does not exist, whereas the rest point S exists. From the inequality 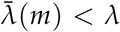, the rest point Q does not exist, so that in this region S is the only non-trivial rest point.

### Characterization of the dynamics

For all the five regions that we identified in the preceding section, Section Stability analysis of the non-trivial rest points provides us with all the information required to determine the stability properties of whichever non-trivial rest points exist. Specifically, when they exist: (i) Q is a sink, (ii) R is a saddle (repelling for points along the TP-edge, attracting for interior points), and (iii) S is a saddle (attracting for points along the HP-edge, repelling for interior points). The stability properties of the trivial rest points T, H, and P in each of the regions are easily identified from Section Dynamics on the edges by using the inequalities defining the five regions. For instance, in the first of the above regions T is a saddle (attracting from H and repelling from P), P is a sink, and H is a source. Together with the fact that there are no interior rest points, we thus obtain the following characterization of the rest points:

i. If (a) *m* ≤ *<u>m</u>* or (b) *m* > *<u>m</u>* and *λ*\*(*m*) < *λ*, then there is no rest point on the edges, T is a saddle (attracting from H and repelling from P), P is a sink, and H is a source. In particular, P is the only stable rest point.
ii. If *m* > *<u>m</u>* and 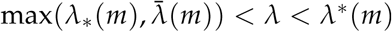, then R is the only rest point on an edge, T is a sink, P is a sink, and H is a source. In particular, T and P are the only stable rest points.
iii. If *m* > *<u>m</u>* and 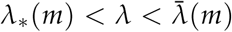, then R and Q are the only rest points on the edges, T is a saddle (attracting from P and repelling from H), P is a sink, and H is a source. In particular, P and Q are the only stable rest points.
iv. If *m* > *<u>m</u>* and 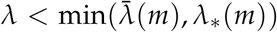, then S and Q are the only rest points on the edges, T is a saddle (attracting from P, repelling from H), and P and H are sources. In particular, Q is the only stable rest point.
v. If *m* > *<u>m</u>* and 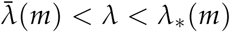, then S is the only rest point on an edge, T is a sink, and P and H are sources. In particular, T is the only stable rest point.

As there are no closed orbits, we further have that the dynamic always (i.e., from all initial conditions) converges to one of the rest points, justifying our focus on the set of stable rest points.

### Characterization of the dynamical regions in the main text

Setting *λ* = *λ*\* (*m*) and solving for *m* we find the critical value

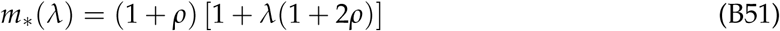

Similarly, setting 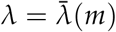 and solving for *m* we find the critical value

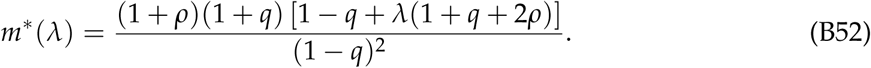

Thus, the curves described by *λ*\*(*m*) and 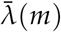 can be equivalently represented by the functions *m*_*_(*λ*) and *m*\*(*λ*). Using this representation, the five dynamical regions identified above then correspond to:

i. *m* < *m*_*_(*λ*).
ii. *λ* > *λ*_*_(*m*) and *m*_*_(*λ*) < *m* < *m*\*(*λ*).
iii. *λ* > *λ*_*_(*m*) and *m* > *m*\*(*λ*).
iv. *λ* < *λ*_*_(*m*) and *m* > *m*\*(*λ*).
v. *λ* < *λ*_*_(*m*) and *m*_*_(*λ*) < *m* < *m*\*(*λ*).

This is the characterization of the dynamical regions that we refer to in the main text, where we label regions *i* to *v* respectively as *P, P+T, P+Q, Q*, and *T*.

### Effects of varying *q* and *ρ*

#### Effects of varying *q*

The critical encounter rate *m*\* is increasing in *q*. Indeed, differentiating equation (B52) with respect to *q* and simplifying, we obtain

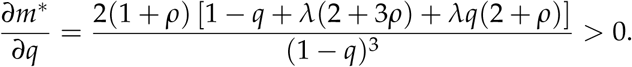

#### Effects of varying *ρ*

The critical availability *λ*_*_ (equation (B46a)) is decreasing in *ρ*. Indeed, differentiating *λ*_*_ (equation (B46a)) with respect to *ρ* and simplifying we obtain

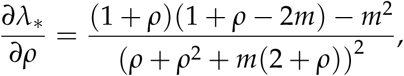

which for *m* > 1 + *ρ* (and hence for *m* > *m*\*) leads to *∂λ*_*_/*∂ρ* < 0.

Both critical encounter rates *m*_*_ (equation (B51)) and *m*\* (equation (B52)) are increasing in *ρ*. Indeed,

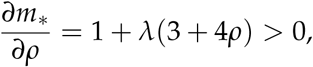

and

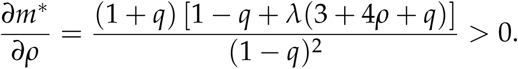

## Appendix C: Alternative Model with Probabilistic Withholding

Here, we consider an alternative model with only two distinct mating strategies: T (”traders”) and N (”non-traders”). The behavior of traders is identical to the one we have considered in our main model. When encountering a potential mate, non-traders provide their eggs with probability *s* and withhold them with probability 1 − *s*. Hence, for *s* = 1 non-traders are equivalent to the providers in our main model, and for *s* = 0 they are equivalent to the withholders that we have considered before.

Our purpose in studying this model is to verify that our conclusions about the conditions under which traders are able to invade a population of non-traders do not qualitatively change when the probability of withholding is determined by the quantitative trait *s* rather than the proportion of withholders among non-trading strategies. To compare like with like, we will consider the condition for a rare mutant trader to invade a resident population of non-traders that provide their eggs with probability *s*\*, where *s*\* is the end-point of the long-term evolution of *s* as predicted by a standard adaptive dynamics analysis (Doebeli, 2011). We will then proceed to compare this invasion condition with the condition under which rare traders can invade the end-point of the evolutionary dynamics along the HP-edge in our main model.

We begin by considering a given, arbitrary *s* and obtain the fitness expressions that will be required for the invasion analysis.

### Female and male reproductive success

Reusing our notation, with *z* now referring to the frequency of non-traders in the population (and *y* = 0), we can write the following expressions for the reproductive success of traders and non-traders in the two sex roles when carrying eggs or not carrying eggs as

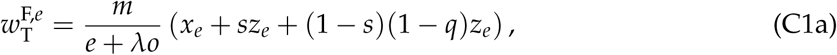

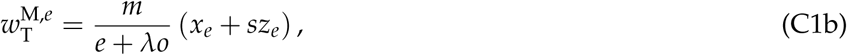

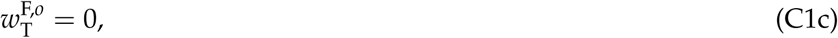

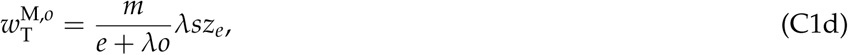

and

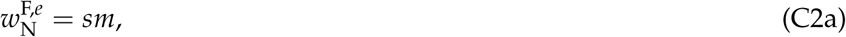

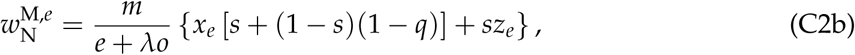

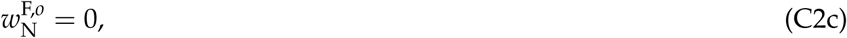

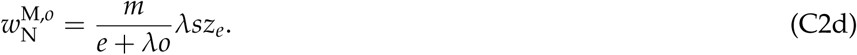

### Demographic dynamics and equilibrium

The demographic dynamics are given by

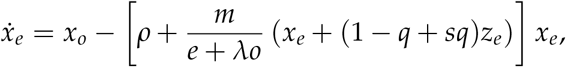

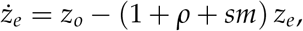

which lead to demographic equilibria satisfying:

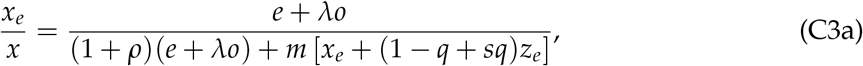

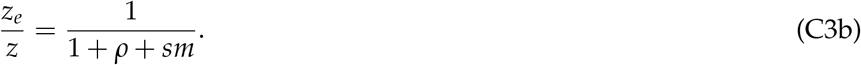

### Expected total reproductive success for each strategy at the demographic equilibrium

The expected total reproductive success for traders is given by

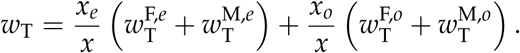

Since *x*_*o*_ = *x* − *x*_*e*_ and, from (C1c), 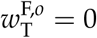, we can write

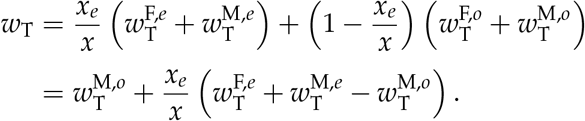

Similarly, for non-traders:

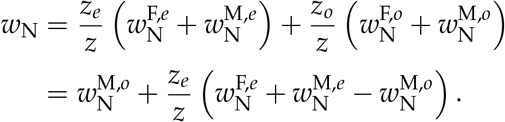

As in the analysis of our main model, we can subtract the common term 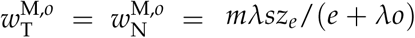, where the equalities are from equations (C1d) and (C2d), and multiply the resulting expressions by *m*/(*e* + *λo*) to obtain renormalized fitnesses. Upon substituting from equations (C1a), (C1b), (C2a), and (C2b), these are:

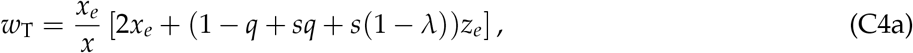

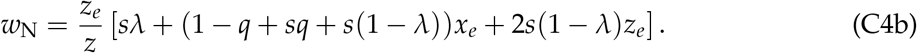

### Adaptive dynamics of the probability of providing in the absence of traders

Consider a resident population of non-traders playing *s* and a small mutant population of non-traders playing *s*′. Let us denote by *z* (resp. *z*′) the proportion of residents (resp. mutants) in the population, and by *z*_*e*_ and *z*_*o*_ (resp. 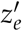 and 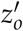) the proportions of residents (resp. mutants) carrying eggs and not carrying eggs. We have 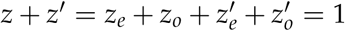 and will assume that *z*′ ≈ 0 (i.e., mutants are rare).

The demographic dynamics of this model are given by

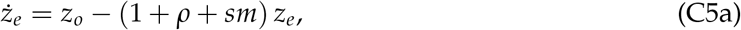

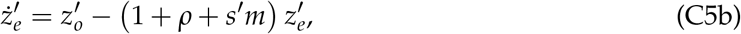

which lead to demographic equilibria given by

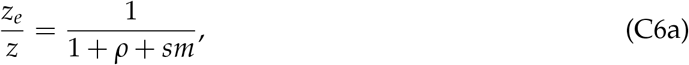

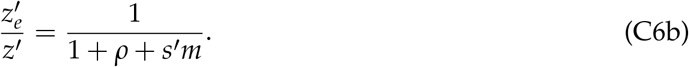

The expressions for female and male reproductive success for the residents, assuming that mutants are so rare that residents only meet resident potential mates are

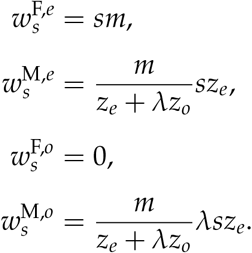

Likewise, for mutants we have

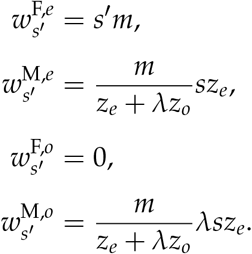

The total expected reproductive success for mutants is given by

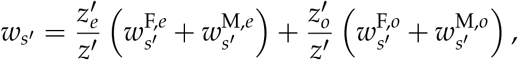

which simplifies to

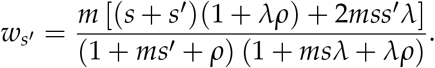

To make progress, we follow the usual assumption in adaptive dynamics that the mutant phenotype *s′* is very close to the resident phenotype *s*. The direction of evolution is then given by the sign of the selection gradient 𝒮(*s*), which can be calculated as

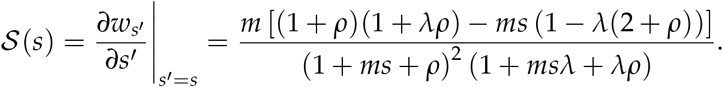

Two different cases arise from this expression. First, the case where *λ* ≥ *λ*_*_ holds, where *λ*_*_ coincides with the critical value for the mating availability we found in our main model (equation (2)). In this case 𝒮(*s*) is positive for all *s ∈* [0, 1). Hence, *s* keeps increasing in evolutionary time until it hits the boundary *s* = 1 so that, for any initial resident population *s*, in the long run non-traders effectively become pure providers (*s*\* = 1). Second, the case in which *λ* < *λ*_*_ holds. In this case the direction of evolution changes sign at a singular interior point *s*\* satisfying 𝒮 (*s*\*) = 0, and therefore given by

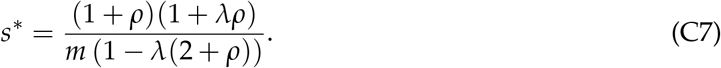

This singular point is convergence stable (i.e., selection drives the evolution of *s* towards this value), as 𝒮 (*s*) changes sign from positive to negative at *s* = *s*\*. Technically, *s*\* is a convergence stable strategy but not an evolutionarily stable strategy (i.e., uninvadable by close mutant trait values), 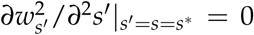. The model can be however adjusted by adding a sufficiently small convex cost to providing (or withholding) eggs so that *s*\* becomes evolutionarily stable without changing the general features of the model.

This general picture corresponds to the one arising from the replicator dynamics model for the competition between providers and withholders (Section HP-edge of Appendix B: Analysis of the Evolutionary Dynamics). Our replicator dynamics model predicts that there can be a polymorphism of withholders and providers (along the HP edge) if and only if *λ* < *λ*_*_ and that otherwise providers dominate withholders. The adaptive dynamics model predicts that there is an interior convergence stable point if and only if *λ* < *λ*_*_ and that otherwise *s*\* = 1. The only important difference in the results is in the expressions for the proportion of providing at equilibrium. In the replicator dynamics model this is given by the value *z*_S_ from (B25), whereas it is given by the value *s*\* in equation (C7) in the adaptive dynamics model. Straightforward algebra shows that *z*_S_ > *s*\* always holds for the relevant parameter values (i.e., when *λ* < *λ*_*_), so that the proportion of providers at the equilibrium in the replicator dynamics model is always less than the convergence stable probability of providing in the adaptive dynamics model.

### Invasion condition for traders

Consider a resident population of non-traders playing *s* = *s*\*. Under which conditions will rare trader mutants invade such a population?

From the analysis above, if *λ* ≥ *λ*_*_ holds, then *s*\* = 1 and non-traders behave as pure providers. Hence, for this parameter constellation the replicator dynamics of traders and non-traders is equivalent to that of traders and providers in the main model (Section TP-edge of Appendix B: Analysis of the Evolutionary Dynamics) and we obtain the equivalent conclusion: if *λ* ≥ *λ*_*_, then rare trader mutants can never invade a monomorphic population of non-traders.

Otherwise, if *λ* < *λ*_*_, then *s*\* is an interior point given by (C7). Subtracting (C4b) from (C4a) and evaluating the resulting expression at *s* = *s*\* and *z* = 1 (which implies *x* = 0 = *x*_*e*_ = *x*_*o*_ = 0) at the demographic equilibrium given by (C3) we get

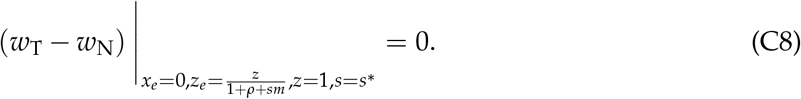

This result is the natural counterpart to the one we obtained in the replicator dynamics model, where the fitness of all three strategies is the same at the rest point S.

In light of (C8), we need to determine the sign of

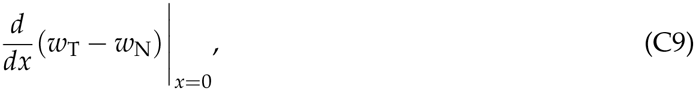

corresponding to the sign of the first-order approximation obtained from the Taylor expansion of the fitness difference *w*_T_ − *w*_N_ around *x* = 0 (*z* = 1), to determine whether or not traders can invade.

Expression (C9) in turn depends on 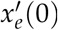, i.e., the derivative of *x*_*e*_ with respect to *x* evaluated at *x* = 0. Letting *z*_*e*_ = (1 − *x*)/(1 + *ρ* + *sm*) in (C3a), implicitly differentiating with respect to *x*, evaluating at *x* = 0, and solving for 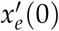, we find

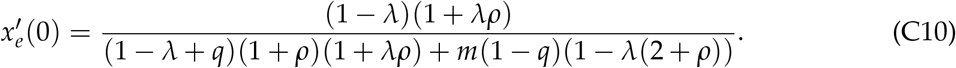

Calculating (C9), substituting (C10), and simplifying, we obtain

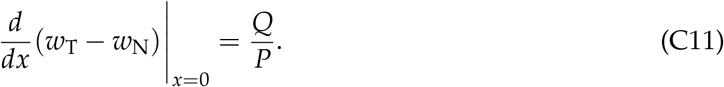

Here,

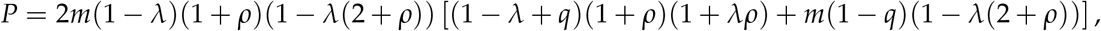

and

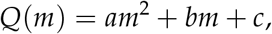

where *a, b*, and *c* are polynomials in *q, λ*, and *ρ*, given by

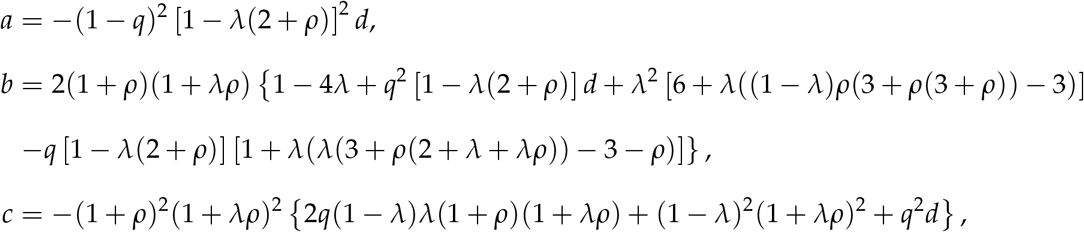

with

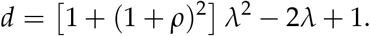

For the relevant parameter constellation (i.e., 1 − *λ*(2 + *ρ*) > 0, so that *s*\* *∈* (0, 1) holds), *P* is always positive. Further, the coefficients *a* and *c* are easily shown to be negative, so that both *Q*(0) < 0 and lim_*m*→∞_ *Q*(*m*) < 0 hold. Hence, the expression in (C11) is negative for such *m*, meaning that traders cannot invade non-traders playing *s*\* if the encounter rate *m* is either sufficiently low or sufficiently high.

A sufficient condition for there to be an intermediate *m* allowing traders to invade is that

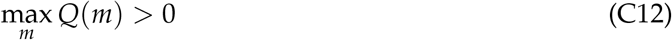

holds. Calculating max_*m*_ *Q*(*m*), we find that a sufficient condition for (C12) to hold is that

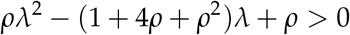

holds, which is equivalent to requiring that 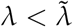 holds, where

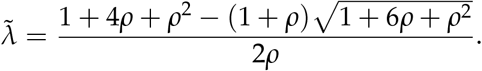

Since 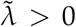 holds for all *ρ* > 0, we conclude that traders can invade a population of non-traders playing *s*\* if the encounter rate is intermediate and if mating availability is sufficiently low. This result is in qualitative agreement with the one we obtained from our main model.

